# Frontal cortex learns to add evidence across modalities

**DOI:** 10.1101/2021.04.26.441250

**Authors:** Philip Coen, Timothy P.H. Sit, Miles J Wells, Matteo Carandini, Kenneth D Harris

## Abstract

To make accurate perceptual decisions, the brain often combines information across sensory modalities. For instance, localizing objects by integrating their image and sound. However, the cortical substrates underlying this audiovisual integration remain uncertain. Here, we show that mouse frontal cortex combines auditory and visual evidence; that this combination is additive, mirroring behavior; and that it evolves with learning. Scanning optogenetic inactivation demonstrated that inactivating frontal cortex impaired choices based on either sensory modality. Recordings from >10,000 neurons indicated that after task learning, activity in frontal area MOs (secondary motor cortex) encodes an additive combination of visual and auditory signals, consistent with the mice’s behavioral strategy. An accumulator model applied to these sensory representations reproduced both the observed choices and reaction times. These results indicate that frontal cortex adapts through learning to combine evidence across sensory cortices, providing a signal that is transformed into a binary decision by a downstream accumulator.

## Introduction

The ability to combine visual and auditory signals to better localize objects in our environment is critical to many species. A simple strategy to combine these signals is to add functions summarizing the evidence from each modality, which is probabilistically optimal if the sources are independent of each other (Gold and Shadlen, 2002; Ma et al., 2006). For instance, the log odds of a stimulus being on the right or left (R vs. L), given independent visual evidence *V* and auditory evidence *A*, is a sum of functions depending only on each modality (Appendix 1):

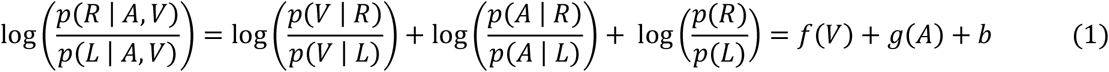

Here, the functions *f*(*V*) and *g*(*A*) quantifying the evidence from each modality may be non-linear, but they are added together linearly. Consistent with this additive strategy, multisensory integration in human and animal behavior is often additive (Alais and Burr, 2004; Angelaki et al., 2009; Chandrasekaran, 2017; Ernst and Banks, 2002; Fetsch et al., 2012; Franklin and Wolpert, 2011; Gu et al., 2008; Körding and Wolpert, 2004; Raposo et al., 2012; Sheppard et al., 2013; Witten and Knudsen, 2005); furthermore, additive evidence combination can be a good heuristic strategy even when the independence assumption is approximate or cannot be verified (Gardner, 2019). Nevertheless, some studies suggest that humans (Battaglia et al., 2003; Colavita, 1974; Sinnett et al., 2007), other primates (Fetsch et al., 2009), and mice (Song et al., 2017) can break this additive law. One specific way that the additive law could be broken is if one modality is dominant, meaning that if sensory information from the modalities conflict, the non-dominant modality is ignored (Song et al., 2017).

The brain could use different strategies to process multisensory signals. One possibility is that there exist specialized regions involved in multisensory, but not unisensory processing: for example, it might be that visual and auditory cortices are necessary and sufficient for responding to visual and auditory stimuli, with a third region required for multisensory processing but having no role in generating behavioral responses to either modality alone. An alternative possibility is that multisensory processing is simply a form of evidence integration: that regions involved in multisensory behavior contain neurons encoding information from both sensory modalities, and have a causal role in behavioral responses to both modalities, alone or in combination. If activity in these regions integrated evidence from the two modalities additively, its output could then drive multisensory behavior according to the additive law (1). In rodents and other mammals, including humans, several brain regions appear to encode multiple modalities, including superior colliculus (Costa et al., 2016; Gharaei et al., 2018; King and Palmer, 1985; Meredith and Stein, 1986; Stein and Stanford, 2008), thalamus (Bieler et al., 2018; Chou et al., 2020; Komura et al., 2005; Lohse et al., 2020), parietal cortex (Angelaki et al., 2009; Avillac et al., 2005, 2007; Fetsch et al., 2012; Gu et al., 2008; Khandhadia et al., 2021; Lippert et al., 2013; Nikbakht et al., 2018; Ohshiro et al., 2011; Raposo et al., 2014; Rohe and Noppeney, 2016; Seilheimer et al., 2013; Song et al., 2017), frontal cortex (Cao et al., 2019; Gu et al., 2015; Jones and Powell, 1970), and even primary sensory cortices (Atilgan et al., 2018; Bizley and King, 2008, 2009; Driver and Noesselt, 2008; Ghazanfar and Schroeder, 2006; Ibrahim et al., 2016; Iurilli et al., 2012; Kayser et al., 2008; Meijer et al., 2017; Meredith and Allman, 2015). However, the causal role of these regions in multisensory decisions remains unclear. Perturbation studies have focused primarily on parietal cortex (Gu et al., 2012; Licata et al., 2017; Raposo et al., 2014) but, with one exception (Song et al., 2017), have not supported a critical role for parietal cortex in multisensory behavior. It is thus not clear which cortical areas, if any, support multisensory decisions, and what multisensory signals are carried by neurons in these regions (Bizley et al., 2016).

## Results

To address these questions, we developed a two-alternative forced choice audiovisual spatial localization task for mice (Figure 1a). We took an established visual detection task where mice turn a steering wheel with their forepaws to indicate the position of a visual grating (Burgess et al., 2017), and extended it by adding a semi-circular array of speakers. On each trial, mice were presented with a flashing visual grating on the left or right screen with variable contrast (including zero contrast, i.e. a grey screen), and a synchronous amplitude-modulated noise played from the left, center, or right speaker. On coherent multisensory trials (auditory and visual stimuli on the same side), and on unisensory trials (either zero visual contrast or central auditory stimulus), mice earned a water reward for indicating the side where the stimuli were presented. On conflict multisensory trials (auditory and visual stimuli on opposite sides), or neutral trials (central auditory and zero contrast visual), mice were rewarded randomly (Figure S1a). On all multisensory trials, auditory and visual stimuli were temporally synchronous. Mice learned to perform this task proficiently (Figure S1b), reaching 96% ± 3% correct (mean ± s.d., n = 17 mice) for the easiest stimuli (coherent trials with the highest visual contrast).

**Figure 1.**
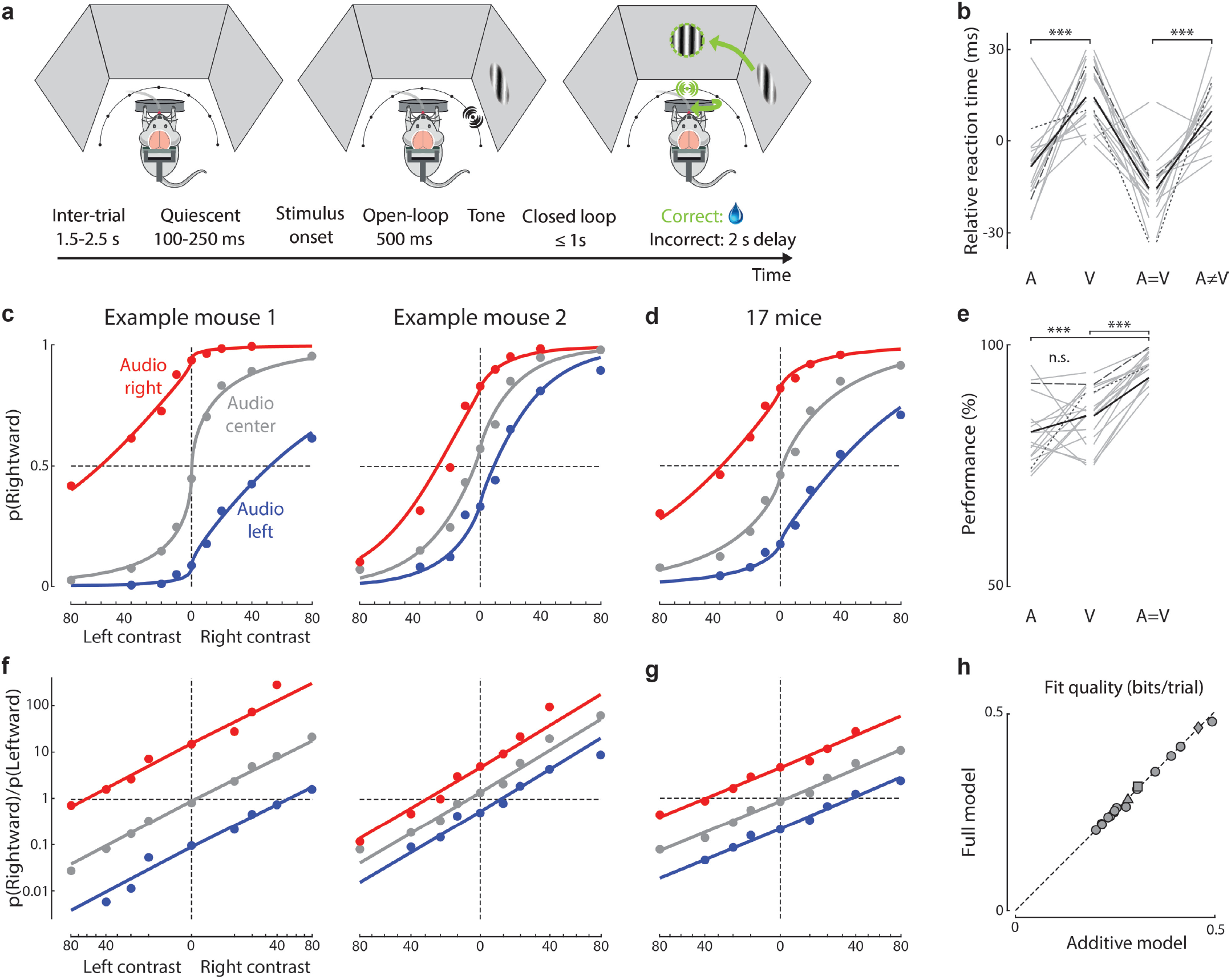
Spatial localization task reveals additive audiovisual integration. **(a)** Behavioral task. Visual and auditory stimuli are presented using 3 screens and an array of 7 speakers. In the example shown, auditory and visual stimuli are both presented on the right, and the subject is rewarded for turning the wheel anti-clockwise to bring the stimuli to the center (a “rightward choice”). Bottom: Task timeline. After an inter-trial interval of 1.5-2.5 s, there is a 100-250 ms quiescent period where mice must hold the wheel still before stimuli appear. They then have 1.5 s to indicate their choice. During the first 500 ms of this period, the stimulus does not move when the wheel is turned (“open loop”), but during the final 1 s stimulus position is yoked to wheel movements. After training, over 90% of choices occurred during open loop period (Figure S1d-f). **(b)** Median reaction times for each stimulus type, relative to the mouse’s mean reaction time across all stimulus types. Only trials with 40% visual contrast were included. Grey lines: individual mice; black line: mean across 17 mice. Long and short dashes indicate example mice from the left and right of (c). ***: p < 0.001 (paired t-test, only comparisons between auditory and visual unisensory, and between conflict and coherent are shown). **(c)** The fraction of rightward choices at each visual contrast and auditory stimulus location for two example mice. Curves: fit of the additive model. **(d)** As in (c), but averaged across 17 mice (~156K trials). Curves: combined fit across all mice. **(e)** Mouse performance (% rewarded trials) for different stimulus types (correct performance on conflict trials is undefined). Plotted as in (b). ***: p < 0.001 (paired t-test), n.s.: p > 0.05. **(f,g)** Same data and fits as c,d, replotted as the odds of choosing right vs. left (in log coordinates, Y-axis), as a function of visual contrast raised to the power γ. In this space, the predictions of the model are straight lines. **(h)** Log_2_-likelihood ratio for the additive model versus an unconstrained model where each combination of visual and auditory stimuli is allowed its own behavioral response that need not follow an additive law. (Both models assessed by 5-fold cross-validation relative to a bias-only model). Triangles and diamonds indicate left and right example mice from (c), squares indicate combined fit across 17 mice. There is no significant difference between the additive and full models (p > 0.05).

Mice performed this task more rapidly when the cues were coherent, suggesting that the task succeeded in engaging a multisensory decision process rather than separate unisensory processes (Figure 1b). Reaction times varied across mice, with a median of 190 ± 120 ms (median ± m.a.d., n = 156k trials in 17 mice; Figure S1h-j), and varied smoothly with stimulus contrast (Figure S1m). In unisensory auditory trials, reaction times were 22 ± 20 ms faster than in unisensory visual trials (s.d., n = 17 mice, p < 0.001, paired t-test), suggesting that the circuits responsible for audiovisual decisions receive auditory spatial signals earlier than visual signals (Hammond-Kenny et al., 2017; Meijer et al., 2018) (Figure 1b, Figure S1o,q). In multisensory trials, reaction times were 25 ± 18 ms faster for coherent than conflict trials (p < 0.001, paired t-test). This acceleration is consistent with a model in which multisensory inputs feed into a single integrator, rather than a “race” model in which two unisensory integrators independently accumulate evidence, with the first to reach threshold producing the decision (Chandrasekaran, 2017; Raab, 1962).

### Spatial localization task reveals additive audiovisual integration

Mice used both modalities to perform the task, even when the two were in conflict. The fraction of rightward choices depended gradually on visual stimulus contrast and markedly increased when sounds were presented on the right and decreased when they were presented on the left (Figure 1c-d, red vs. blue). Mice performed more accurately on coherent trials than unisensory trials of either modality (Figure 1e, Figure S1n,p), indicating that they were attending to both modalities (Alais and Burr, 2004; Hammond-Kenny et al., 2017; Meijer et al., 2018; Pisupati et al., 2021; Raposo et al., 2012; Stein et al., 1989).

To test whether mice make multisensory decisions according to the additive law, we fit an additive model to their behavioral responses. *Eqn*. 1 can be rewritten as:

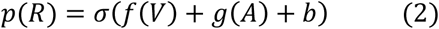

where *p*(*R*) is the probability of making a rightward choice, and *σ*(*x*) = 1/(1 + *exp*(−*x*)) is the logistic function. We first fit this additive model in a nonparametric manner (i.e. with no constraints on the functions *f* and *g*) and found that it provided excellent fits (Figure S2f). We then further simplified it by modelling *f* parametrically, using a power function to account for contrast saturation in the visual system (Zatka-Haas et al., 2021):

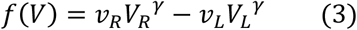

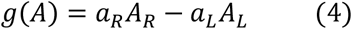

Here *V_R_* and *V_L_* are right and left visual contrasts (at least one of which was always zero), and *A_R_* and *A_L_* are indicator variables for right and left auditory stimuli (with value 1 or 0 depending on the auditory stimulus position). This additive parametric model gave performance almost as good as the 11-parameter nonparametric additive model (Figure S2f) but has only 6 free parameters: the bias *b*, the visual exponent *γ*, two visual sensitivities *v_R_* and *v_L_*, and two auditory sensitivities *a_R_* and *a_L_*. From here on, we refer to this additive parametric model simply as the “additive model”.

The additive model provided excellent fits to the multisensory decisions of all the mice. It fit the choices of individual mice (Figure 1c) as well as the mean choice averaged across mice (Figure 1d). A particularly clear view of the model can be obtained by replotting these data in terms of log odds of rightward vs. leftward choices (as in *Eqn*. 1) vs. linearized visual contrast (contrast raised by the exponent *γ, Eqn.3*). In this representation, the responses to unisensory visual stimuli fall on a line, and the effect of adding auditory cues is to shift this line additively (Figure 1f,g, Figure S3a-o). The intercept of the line is determined by the bias *b*, its slope by the visual sensitivity *v*, and the additive offset by the auditory sensitivity *a*.

The additive model performed better than non-additive models where one modality dominates the other during conflicts or models of sensory bias (Figure S2a-e). Indeed, it performed as well as a full model, which used 25 parameters to fit the response to each stimulus combination and was not constrained to be additive (Figure 1h). The additive model also captured the results of experiments where we tested additional auditory conditions (Figure S1k-l), and it could be fit from the responses obtained in the unisensory trials alone (Figure S3p). As predicted by the additive model, equal and opposite auditory and visual stimuli (i.e. stimuli eliciting an equal probability of moving in opposite directions when presented alone) led to neutral behavior when presented together, i.e. a 50% chance of moving left or right (Figure S2g). In contrast, a model of sensory dominance would predict that equal and opposite stimuli would lead to consistent movements in the direction suggested by the stimulus of dominant modality.

### Optogenetic inactivation identifies roles of sensory and frontal cortical areas

To determine which regions of dorsal cortex are necessary to perform the task, we used laser-scanning photo-inactivation. To identify regions potentially involved in integrating auditory and visual evidence, we looked for sites where inactivation impaired choices on unisensory trials of either modality, as well as multisensory trials.

We optogenetically inactivated 52 cortical sites, by transcranial laser illumination in mice expressing ChR2 in parvalbumin interneurons (Cardin et al., 2009; Guo et al., 2014; Olsen et al., 2012; Zatka-Haas et al., 2021) (3 mW; 462 nm; 1.5 s duration following stimulus onset; Figure 2a). We combined results across mice and hemispheres because inactivation effects were qualitatively consistent and symmetric (Figure S4a-b). Control measurements established that mouse choices were not affected when we targeted locations on the dental cement outside the brain (Figure S4c). Due to light scattering in the brain, we expect inactivation to significantly impact areas ~1 mm from the target location at our laser power (Hao et al., 2021; Li et al., 2019; Zatka-Haas et al., 2021) (Figure 2a). For this reason, and because brain curvature hides auditory cortex, laser sites on “lateral sensory cortex” between primary visual and auditory areas likely inactivated both visual and auditory cortices. We observed distinct impacts on task performance by inactivation in three cortical regions: visual, lateral sensory, and frontal (Figure 2b-e).

**Figure 2.**
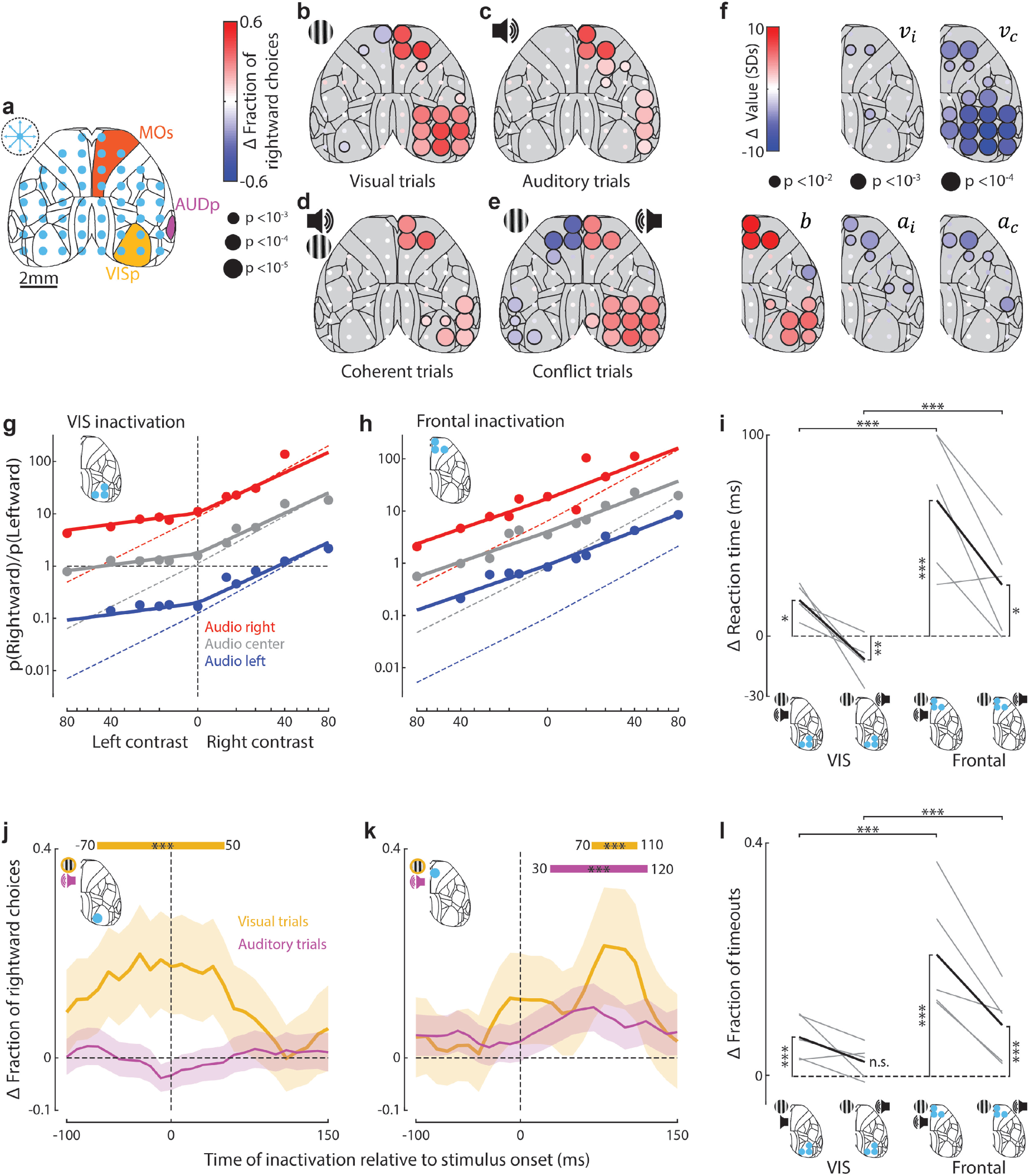
Optogenetic inactivation identifies roles of sensory and frontal cortical areas. **(a)** Schematic of inactivation sites. On ~75% of trials, a blue laser randomly illuminated regions centered on one of 52 locations in either hemisphere (blue dots), from stimulus onset (fixed duration of 1.5 s). Dashed circle indicates estimated size of cortical area inactivated by laser stimulation (1 mm radius). Yellow, orange, and magenta regions indicate primary visual (VISp), primary auditory (AUDp) and secondary motor (MOs) regions for reference. Sites that inactivated the dorsal aspect of auditory cortex also inactivated lateral visual cortex. **(b)** Change in the fraction of rightward choices on inactivation of 52 laser sites for unisensory left visual stimulus trials. Dot color indicates change in the fraction of rightward choices; dot size represents statistical significance (5 mice, permutation test, see Methods). Data for right stimulus trials were also included in the average after reflecting the maps (see Figure S4a for both individually). **(c)** As in (b), but for unisensory auditory trials. **(d)** As in (b), but for coherent multisensory trials. **(e)** As in (b), but for conflict multisensory trials. **(f)** As in (b-e), but dot color indicates the change in parameters of the additive model. Here, the bias b represents bias toward ipsilateral choices (relative to the site of inactivation); v_i_ and v_c_ represent sensitivity to ipsilateral and contralateral visual contrast; α_i_ and α_c_ represent sensitivity to ipsilateral and contralateral auditory stimuli. **(g)** Fit of the additive model to trials when right visual cortex was inactivated, combined across the three sites shown in the inset. Dashed lines indicate model fit to non-inactivation trials. Data combined across 5 mice. Trials with inactivation of left visual cortex were also included in the average after reflecting the maps (5 mice, 6497 trials). The effect of inactivation on model parameters was significant (paired t-test across 5 mice, p < 0.05). **(h)** As in (g), but for trials when frontal cortex was inactivated (5 mice, 5612 trials). The effect of inactivation on model parameters was significant (paired t-test across 5 mice, p < 0.05). **(i)** Change in multisensory reaction times when visual or frontal cortex were inactivated contralateral to the visual stimulus. Grey and black lines indicate individual mice (n = 5) and the mean across mice. Reaction times were calculated as the mean across the median for each contrast. 0 indicates the reaction time for non-inactivation trials. Values above 100 ms were truncated to 100 ms for visualization but not analyses. On coherent trials, inactivation of either visual or frontal cortex significantly increased reaction time and the effect was larger for frontal. On conflict trials, inactivation of visual cortex decreased reaction time while inactivation of frontal cortex caused an increase. *: p < 0.05, **: p < 0.01, ***: p < 0.001 (linear mixed effects model). **(j)** The change in fraction of rightward choices when contralateral visual cortex was inactivated on visual (yellow, 519 inactivation trials) or auditory (magenta, 1205 inactivation trials) trials. Inactivation was a 25 ms, 25 mW laser pulse at different time points relative to stimulus onset. Curves show the average pooled over all sessions and mice, smoothed with a 70 ms boxcar window. Shaded areas: 95% binomial confidence intervals. *** indicate the intervals in which the fraction of rightward choices differs significantly from control trials (p < 0.001, Fisher’s exact test). **(k)** As in (j), but for inactivation of frontal cortex (451 and 1291 inactivation trials for auditory and visual stimulus conditions). **(l)** As in (i), but for the change in the fraction of timeout trials. On coherent trials, inactivation of either visual or frontal cortex significantly increased timeouts and the effect was larger for frontal. On conflict trials, only frontal inactivation significantly changed the fraction of timeouts, causing them to increase above control and visual cortex levels. ***: p < 0.001 (linear mixed effects model).

Inactivating visual cortex impaired visual but not auditory choices. As seen in other visual tasks (Burgess et al., 2017; Glickfeld et al., 2013; Zatka-Haas et al., 2021), inactivation of visual cortex reduced responses to contralateral visual stimuli, whether presented alone (Figure 2b) or together with auditory stimuli (Figure 2d,e). It had a smaller effect in coherent trials, when those same choices could be based on audition alone, than in unisensory visual trials (Figure 2d, p < 0.01, paired t-test across 5 mice). Conversely, it did not affect choices based on unisensory auditory stimuli (Figure 2c), indicating that in this task visual cortex does not play a substantial role in processing auditory signals.

Inactivating frontal cortex impaired choices based on either modality, with similar strength (Figure 2b-e). On visual trials, inactivating frontal cortex had a similar effect to inactivating visual cortex: it reduced responses to contralateral stimuli (Figure 2b, as in visual detection tasks (Burgess et al., 2017; Glickfeld et al., 2013; Zatka-Haas et al., 2021)). However, inactivation of frontal cortex also reduced responses to contralateral auditory stimuli to a similar extent (p > 0.05, t-test across mice; Figure 2c). In coherent multisensory trials, inactivation of frontal cortex reduced responses when both stimuli appeared on the contralateral side, but had no significant effect when they both appeared on the ipsilateral side (Figure 2d). On conflict trials, where one stimulus appeared on each side, frontal inactivation reduced the responses to the contralateral stimulus, whichever modality it came from (Figure 2e). These data are thus consistent with a role for frontal cortex in integrating visual and auditory evidence.

Finally, inactivating lateral sensory cortex impaired visual choices strongly, and auditory choices to a weaker extent. It decreased correct responses to contralateral stimuli whether visual alone (Figure 2b), auditory alone (Figure 2c), or combined (Figure 2d,e), but had a larger effect on visual than auditory choices (Figure 2b-c, p < 0.05, t-test across mice). These results might suggest a multisensory role for this lateral region but might more simply arise from light spreading into visual and auditory cortices: indeed, due to brain curvature, light arriving on this region is likely to reach auditory cortex (which is required for auditory localization (Jenkins and Merzenich, 1984)) but only after passing through overlying tissue, which may explain why a weaker effect was seen for auditory stimuli.

The results of these inactivations were well captured by the additive model. The model accounted for the effects of inactivating visual cortex via a decrease in the sensitivity for contralateral visual stimuli *v_c_* (Figure 2f), which reduced performance for contralateral visual stimuli regardless of auditory stimuli (Figure 2g). Inactivating lateral sensory cortex had a similar impact on visual sensitivity *v_c_*, but also more weakly decreased contralateral auditory sensitivity *a_c_* (Figure 2f, Figure S4e, p < 0.07, t-test across mice). Inactivating frontal cortex reduced both visual and auditory sensitivity by a similar amount (p > 0.65, t-test across mice), and also increased bias *b* to favour ipsilateral choices (Figure 2f,h, Figure S4g). The effects of inactivating visual, lateral, and frontal cortices were statistically different to each other (Figure S4h). For example, inactivating frontal cortex reduced sensitivity to both contralateral and ipsilateral stimuli but inactivating lateral sensory cortex only reduced sensitivity to contralateral stimuli (Figure 2f).

The effect of cortical inactivation on reaction times revealed a difference between frontal and other cortices. Inactivating frontal cortex increased reaction times in all stimulus conditions (Figure 2i, Figure S5a-h). In contrast, the effect of inactivating visual cortex depended on the stimulus: responses to contralateral visual stimuli, or coherent contralateral audiovisual stimuli were delayed, but responses to conflicting stimuli with a contralateral visual component were accelerated (Figure 2i, Figure S5a-h), as if inactivation of visual cortex causes the brain to ignore the contralateral visual stimulus, and respond as on unisensory auditory trials (Figure 1b). The effects of inactivating the lateral cortex were similar to visual cortex but did not reach statistical significance. Similar results were seen in the fraction of timeouts, i.e., trials where the mouse failed to respond within 1.5 s: Inactivation of frontal cortex increased timeouts in every stimulus condition, irrespective of modality and laterality, while inactivation of visual and lateral sensory regions only impacted contralateral visual and coherent stimuli, and had a weaker effect (Figure 2l, Figure S5i-p). These data suggest that inactivating visual or lateral cortex has a similar role to the contralateral stimulus being absent, which may speed or slow reaction times depending on whether it resolves a stimulus conflict; but that inactivating frontal cortex impairs a process of multisensory evidence integration, slowing all choices and especially contralateral ones (p < 0.001, linear mixed effects model).

The critical time for inactivation of visual cortex was earlier than that of frontal cortex. We used 25 ms laser pulses to briefly inactivate visual and frontal cortex at different timepoints relative to stimulus onset on unisensory trials (Zatka-Haas et al., 2021) (see Methods). Inactivating right visual cortex significantly increased the fraction of rightward choices from 70 ms prior to 50 ms after the appearance of a visual stimulus on the left (p < 0.001) but had no significant effect at any time after an auditory stimulus (Figure 2j). Conversely, frontal inactivation impacted behavior when contralateral to either visual (between 80 and 110 ms after stimulus) or auditory (between 30 and 120 ms) stimuli (Figure 2k). The earlier critical window for frontal inactivation on auditory trials is consistent with the faster reaction times on these trials (Figure 1b). However, in both cases, inactivation of frontal cortex had no significant effect from 120 ms after stimulus onset, suggesting that after this time, its role in sensory integration is limited. We did not observe any significant effect of these short inactivation pulses when stimuli were ipsilateral to the site of inactivation or when inactivation was targeted outside the brain. Thus, visual cortex’s role in the task is earlier and unisensory, while frontal cortex is later and multisensory.

### Neurons in frontal area MOs encode stimuli and predict behavior

The fact that frontal inactivation impairs decisions based on either visual or auditory stimuli or their combination, suggests that at least some areas of frontal cortex may integrate evidence from both modalities. To test this hypothesis, and to determine which specific parts of frontal cortex are most involved, and what sort of neural code they might use to represent the integrated information, we recorded acutely with Neuropixels probes during behavior (Figure 3a-d). We recorded 14,656 neurons from frontal cortex across 88 probe insertions (56 sessions) from 6 mice (Figure 3a, Figure S6a-b) divided across the following areas: secondary motor (MOs, 3041 neurons), orbitofrontal (ORB, 5112 neurons), anterior cingulate (ACA, 727 neurons), prelimbic (PL, 1,332 neurons), and infralimbic (ILA, 1254 neurons). We also recorded from 2,068 neurons in nearby olfactory areas (OLF). These regions exhibited a variety of neural responses, including neurons that were sensitive to visual and auditory location (Figure 3b-c), and to the animal’s upcoming choice (Figure 3d).

**Figure 3.**
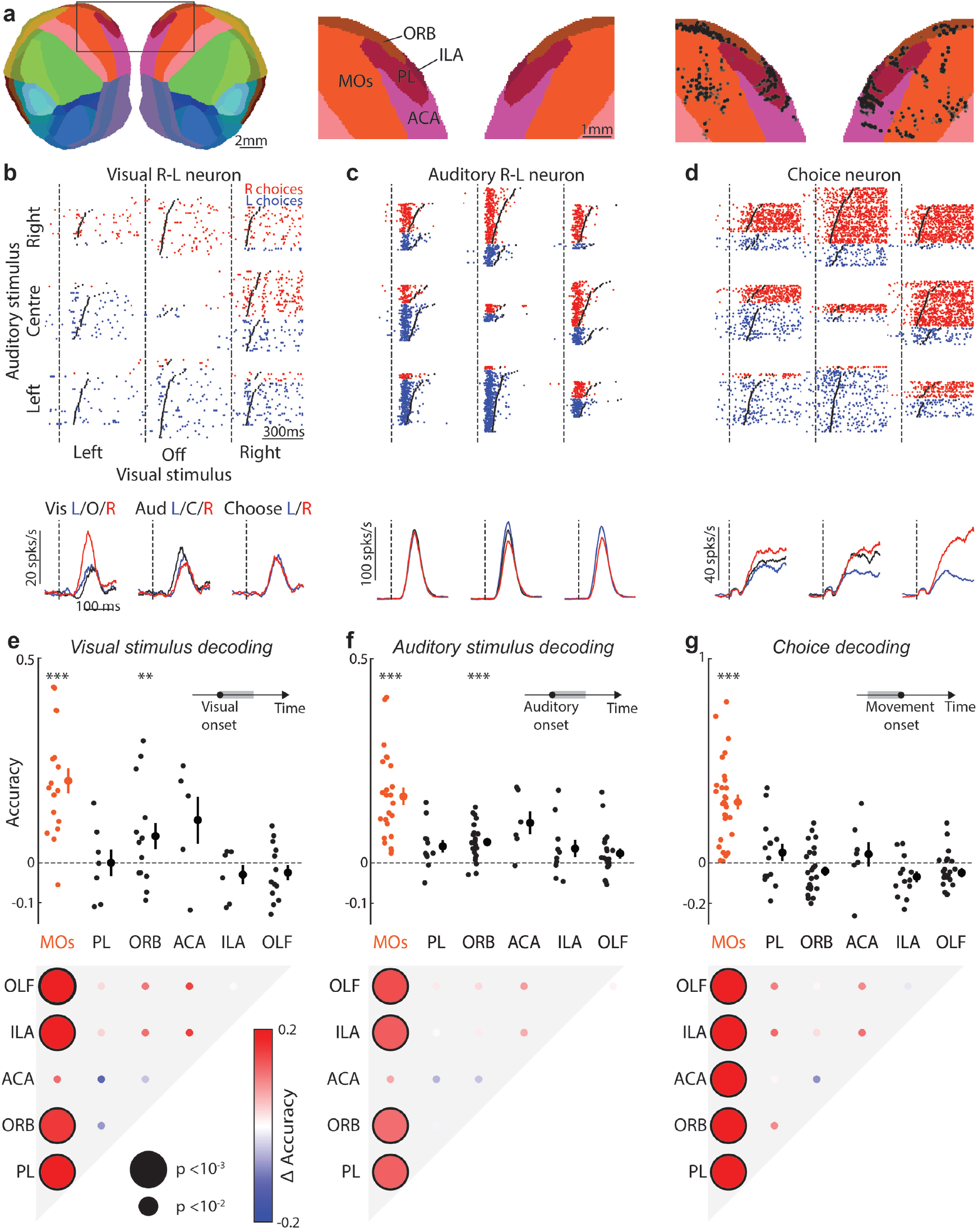
Neurons in frontal area MOs encode stimuli and predict behavior. **(a)** Recording locations for cells (black dots, right), overlayed on a flattened cortical map (using the Allen CCF(Wang et al., 2020)), showing locations in secondary motor (MOs), orbitofrontal (ORB), anterior cingulate (ACA), prelimbic (PL) and infralimbic (ILA) areas. **(b)** Top: Raw spike rasters, separated by trial condition, from a neuron sensitive to visual spatial location (d-Prime = 1.85). Red/blue rasters indicate trials where the mouse made a rightward/leftward choice. Dashed line and black points represent stimulus onset and movement initiation. Bottom: Peri-stimulus time histogram (PSTH) of the neural response, averaged across different visual (left), auditory (center) or choice (right) conditions. Trials are not balanced: choice and stimulus location can be correlated. **(c)** As in (b), for a neuron that is sensitive to auditory spatial location (d-Prime = −0.81). **(d)** As in (b), for a neuron that is sensitive to the animal’s choice (d-Prime = 2.61). **(e) Top:** Cross-validated accuracy (relative to a bias model, see Methods) of an SVM decoder trained to predict visual stimulus location from population spiking activity time-averaged over a window 0 ms to 300 ms after stimulus onset. Accuracies 0 and 1 represent chance and optimal performance. Each point represents the decoding accuracy from neurons in one of the regions labelled in (a), or olfactory areas (OLF), from a single experimental session. For each point, 30 neurons were randomly selected to equalize population size across sessions and brain regions. ***: p < 0.001, **: p < 0.01 (≥ 5 sessions from 2-5 mice for each region, t-test). **Bottom:** Significance for inter-region comparison of decoding accuracy (Linear mixed effects model). Black outlines indicate statistically significant pairwise difference, dot size indicates significance level. **(f)** As in (e), but for decoding of auditory stimulus location (≥ 6 sessions, 3-6 mice). **(g)** As in (e), but for decoding choices based on spiking activity time-averaged over a window 0 ms to 130 ms preceding movement onset (≥ 7 sessions, 3-6 mice).

Of the fontal regions we recorded, task information was represented most strongly in secondary motor area MOs. To identify regions encoding task information, we trained separate linear support vector machine (SVM) decoders to decode stimulus location or upcoming choice from neural activity. MOs was the only region able to predict the animal’s upcoming choice before movement onset (Figure 3g, Figure S6c), and encoded auditory and visual stimulus location significantly more strongly than other recorded regions (Figure 3e-f; p<0.01, Linear mixed effects model; in ACA the difference did not reach significance). We observed consistent results at the single cell level: neurons in MOs yielded higher discriminability for stimulus location and choice than all other regions (Figure S6e-f; p<0.05, Linear mixed effects model; in ACA and PL the difference did not reach significance for visual location). These observations were robust to the correlation between stimuli and choices: even when controlling for this correlation, MOs still had the largest fraction of neurons with significant coding of stimulus location or pre-movement choice (Figure S6g-h). Once movements were underway, however, we could decode their direction from multiple regions, consistent with observations that ongoing movements are encoded throughout the brain (Musall et al., 2019; Stringer et al., 2019) (Figure S6d).

### Frontal area MOs integrates task variables additively

Given the additive effects of visual and auditory signals on behavior, we asked whether these signals also combine additively in MOs neural activity. To test this hypothesis, we used an ANOVA-style decomposition into temporal kernels (Park et al., 2014) to analyze responses to combined audiovisual stimuli during behavior. For simplicity, we focused on trials of a single visual contrast, so that we could define binary variables *a_i_,v_i_,c_i_* = ±1 encoding the laterality (left vs. right) of auditory stimuli, visual stimuli, and choices. We could then decompose ***F**_i_*(*t*), the firing rate on trial *i* at time *t* after stimulus onset, as the sum of 6 temporal kernels:

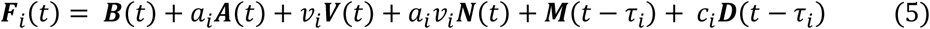

Here, ***B*** is the mean stimulus response averaged across stimuli, ***A*** and ***V*** are the additive main effects of auditory and visual stimulus location, and ***N*** is a non-additive interaction between them. Finally, ***M*** is a kernel for the mean effect of movement (regardless of direction, and relative to *τ_i_*, the time of movement onset on trial *i*) and ***D*** is the differential effect of movement direction (right minus left). To test for additivity, we compared performance of this full model against a restricted additive model where the non-additive interaction term is 0 (*N* = 0), and used cross-validation to compare the two models.

The results were consistent with additive integration of visual and auditory signals for stimulus location. Indeed, the additive neural model outperformed the full model with interactions between visual and auditory stimuli (Figure 4a-c), as well as an alternative full model with interactions between stimuli and movement (Figure S7a). Similar results were seen during passive presentation of task stimuli, when sensory responses could not be confounded by activity related to movement (Figure 4d-f, Figure S7h).

**Figure 4.**
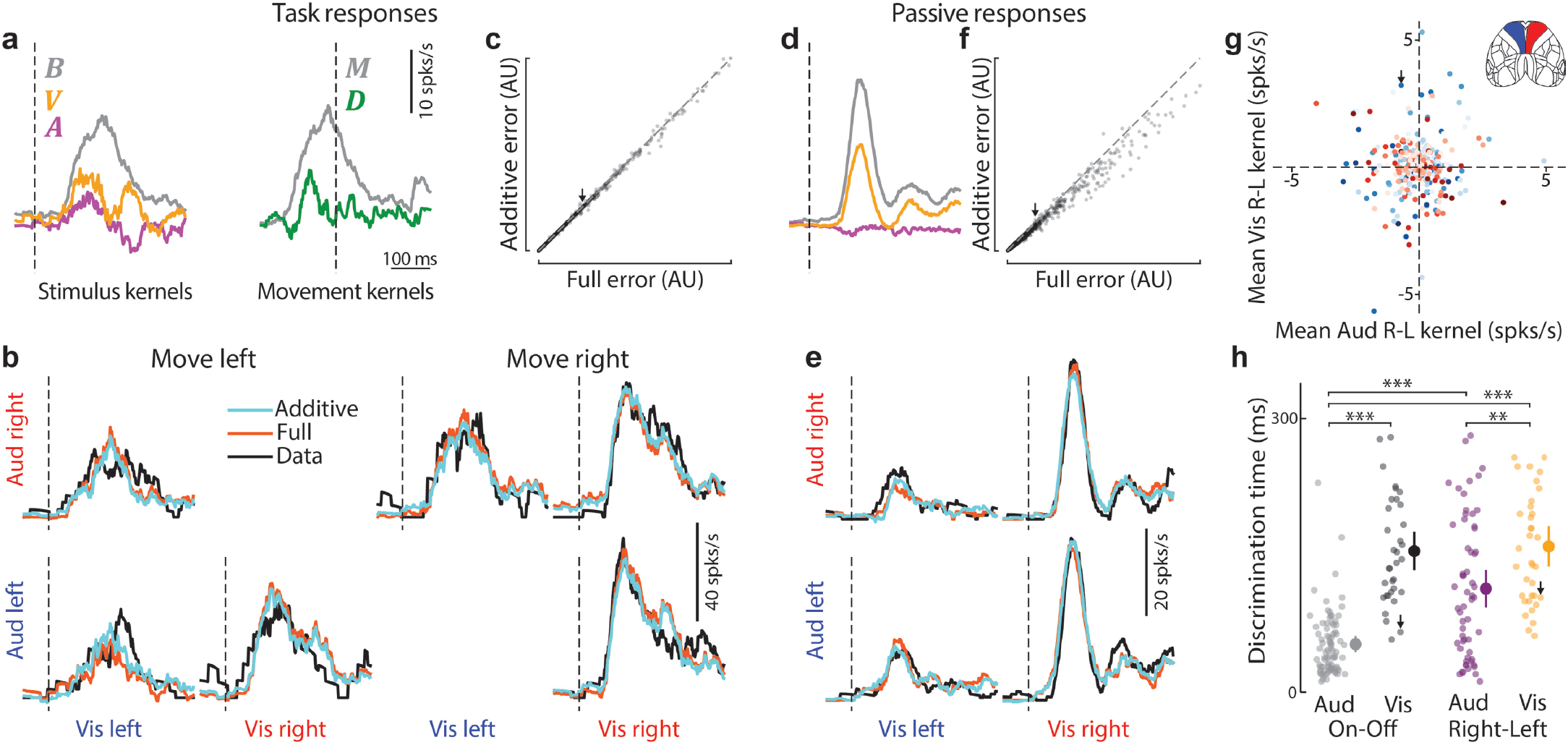
Frontal area MOs encodes task variables additively. **(a)** Example kernels from fitting the additive neural model to an example neuron. Dashed lines indicate stimulus onset for stimulus kernels and movement onset for movement kernels. **B:** overall mean stimulus response, **A** and **V:** additive main effects of auditory and visual stimulus location. **M** is a kernel for the mean effect of movement (regardless of direction, and relative to τ_i_, the time of movement onset on trial i) and D is the differential effect of movement direction (right minus left). To obtain these fits, the non-additive kernel N was set to 0. **(b)** Cross-validated model fits for the example neuron from (a) to neural activity averaged over each combination of stimuli and movement direction. Trials where stimuli were coherent but the mouse responded incorrectly were excluded due to low trial numbers. Cyan and orange lines show predictions of the additive (**N** = 0) and full models, black line shows test-set average responses. Dashed lines indicate stimulus onset. **(c)** Prediction error (see Methods) across all neurons for the additive and full models. Arrow indicates position of the example cell in (a-b). The additive model has smaller prediction error (p = 0.037, linear mixed effects model, 2183 cells, 5 mice). The largest 1% of errors were excluded from plot for visualization, but not from statistical analysis. **(d)** As in (a), but for neural activity during passive stimulus presentation, using only sensory kernels. **(e)** As in (b), but fitting sensory kernels to all audiovisual combinations presented in passive conditions. **(f)** As in (c), but for passive stimulus conditions. The additive model has smaller prediction error. p < 10^-10^ (2509 cells, 5 mice, linear mixed-effects model). **(g)** Encoding of visual stimulus preference (time-averaged amplitude of kernel, V) versus encoding of auditory stimulus preference (time-averaged amplitude of kernel, A) for each cell. There was no correlation between auditory and visual directional preference. p > 0.05 (2509 cells, Pearson correlation test). Red/blue indicates whether the cell was recorded in the right/left hemisphere and saturation indicates fraction of variance explained by the sensory kernels. **(h)** Discrimination time (see Methods) relative to stimulus onset during passive conditions. Auditory Right-Left neurons (sensitive to auditory location, magenta, n = 59) discriminated stimulus location earlier than Visual Right-Left neurons (gold, n = 36). Auditory On-Off neurons (sensitive to auditory presence, but not necessarily location, grey, n = 82) discriminated earlier than all other neuron classes, including Visual On-Off (n = 36, black). Points are individual neurons and bars indicate standard error. **: p < 0.01, ***: p < 0.001 (Mann–Whitney U test).

MOs neurons provided a mixed representation of visual and auditory stimulus locations but encoded the two modalities with different time courses. Consistent with reports of mixed multisensory selectivity in the parietal cortex of rat (Raposo et al., 2014) and primate (Gu et al., 2008), the auditory and visual stimulus preferences of MOs neurons were neither correlated nor lateralized: cells in either hemisphere could represent the location of auditory or visual stimuli with a preference for either location, and could represent the direction of the subsequent movement with a preference for either direction (Figure 4g, Figure S7c-g). Neurons that responded to one modality, however, also tended to respond to the other, as evidenced by a weak correlation in the absolute sizes of the auditory and visual kernels (Figure S7b). Nevertheless, representations of auditory and visual stimuli had different time courses: neurons could distinguish the presence and location of auditory stimuli earlier than for visual stimuli (Figure 4h). This is consistent with the more rapid behavioral reactions to auditory stimuli (Figure 1b) and the earlier critical window for MOs inactivation on unisensory auditory than visual trials (Figure 2j-k). Indeed, the earliest times in which MOs encoded visual or auditory stimuli (Figure 4h) matched the times for which MOs inactivation impacted behavioral performance (Figure 2j-k).

MOs encoded information about auditory onset (regardless of sound location) more strongly and significantly earlier than information about visual onset or the location of either stimulus (Figure 4h, Figure S7i). This may explain why mice in a detection task, rather than our localization task, exhibit auditory dominance in multisensory conflict trials (Song et al., 2017).

### MOs only encodes multisensory signals after task training

The discriminability of both auditory and visual location was higher in neural populations recorded in trained mice (Figure 5a-b). We recorded the responses of 2,702 MOs neurons to the task stimuli in 4 naïve mice and compared their activity to that previously characterized in trained mice. Encoding of visual stimulus location was significantly increased after training (**: p < 0.01, Welch’s t-test), and the discriminability index (d’: the absolute mean difference of firing rates between stimulus conditions divided by the mean of the standard deviations across trials for each stimulus condition) of neurons recorded in naïve mice was not significantly different from a shuffled control (Figure 5a). Furthermore, encoding of auditory stimulus location was weaker in naïve mice (Figure 5b, p < 0.01, Welch’s t-test), though still significant (p < 0.01, Welch’s t-test).

**Figure 5.**
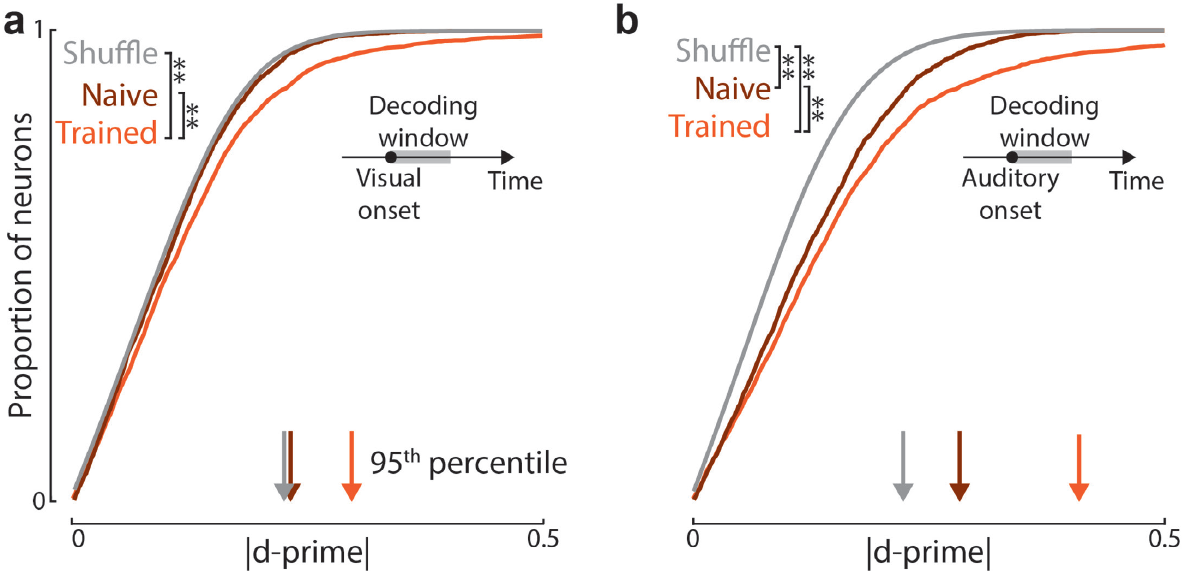
Audiovisual integration in MOs develops through learning. **(a)** Cumulative histogram of absolute discriminability index (d-prime) scores for MOs neurons in response to visual stimuli in naïve mice, trained mice, and shuffled data. The proportion of spatially sensitive neurons is enriched by training (**: *p* < 0.01, Welch’s t-test). Neurons in naïve mice did not have significantly higher discriminability than after shuffling stimulus location (p > 0.05, Welch’s t-test). Arrows indicate the value of d-prime at the 95^th^ percentile for each category. **(b)** As in (a), but for auditory stimuli. The proportion of spatially sensitive neurons is enriched by training. Neurons in naïve mice did have significantly higher discriminability than after shuffling stimulus locations (**: *p* < 0.01, Welch’s t-test).

### An accumulator applied to MOs activity reproduced decisions

We asked whether the representation of multisensory task stimuli in MOs could explain the properties of the animal’s behavioral choices. To establish this, we considered an accumulator model that makes choices based on MOs population activity (Figure 6a,b). To avoid the confound of movement encoding in MOs, we used passive stimulus responses, and generated surrogate population spike trains ***x***(*t*) by selecting (from all recordings) MOs neurons encoding the location of at least one of the sensory stimuli. These spike trains were fed into an accumulator model (akin to a drift diffusion model(Gold and Shadlen, 2002, 2007; Ratcliff, 1978)): they were linearly integrated over time to produce a scalar decision variable *d*(*t*):

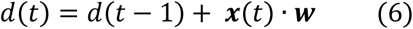

**Figure 6.**
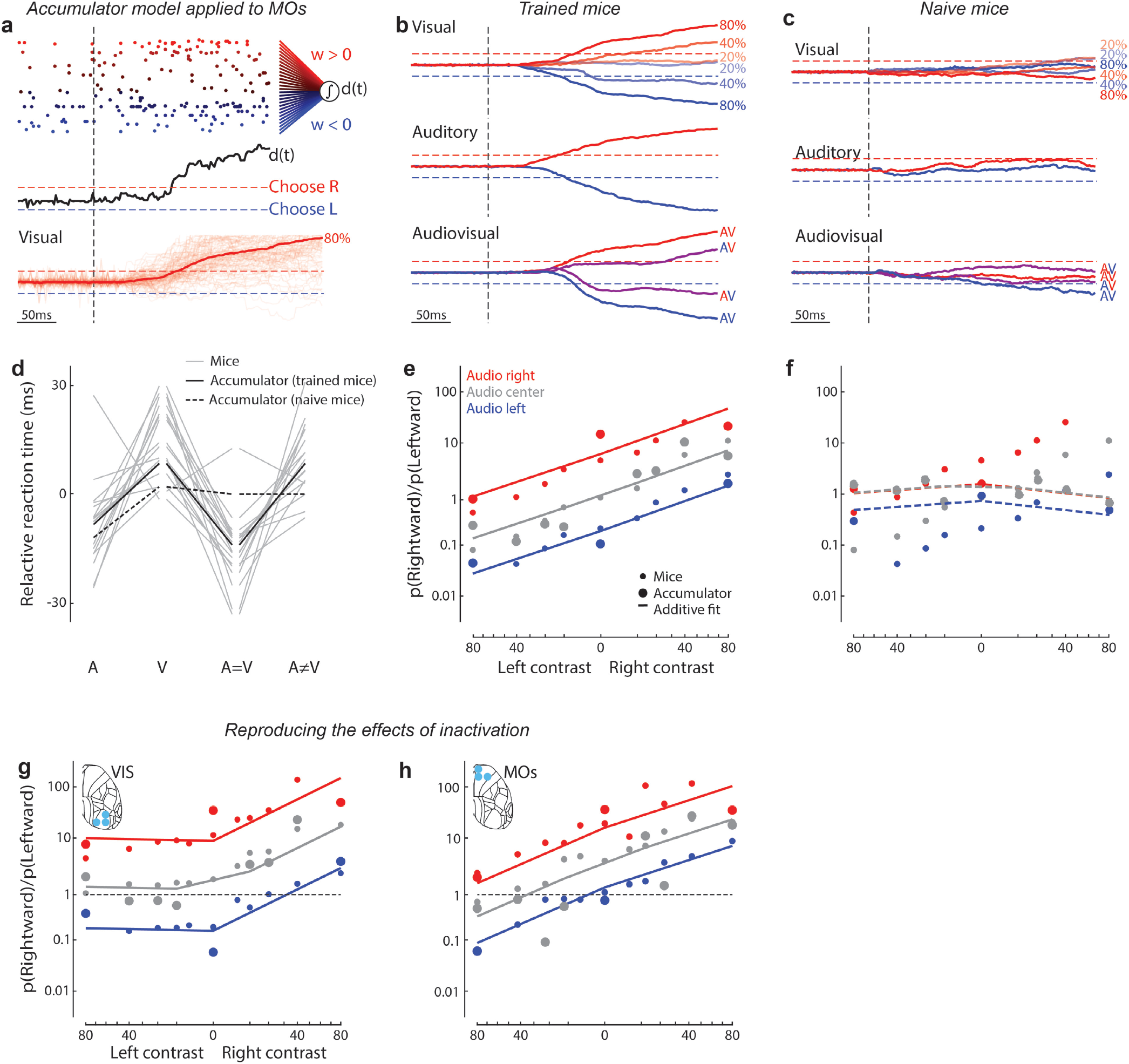
An accumulator applied to MOs activity in trained mice reproduced decisions. **(a) Top**: population spike train rasters for a single trial, colored according to the fitted weight for that neuron, with red and blue indicating that spiking will push the decision variable d_t_ to the rightward or leftward decision boundary. Vertical dashed line represents stimulus onset. Population activity was created from neurons of multiple passive recording sessions in MOs of trained mice. **Middle**: Evolution of the decision variable over this trial. Red/blue dashed lines indicate the rightward/leftward decision boundaries. **Bottom**: Sumperimposed examples of the decision variable trajectory for multiple unisensory visual trials with 80% rightward contrast (thin) and their mean (thick). **(b)** Mean decision variable trajectory for visual-only (top), auditory-only (middle), and multisensory (bottom) stimulus conditions. **(c)** As in (b), but for an accumulator model trained on neural activity created from the same number of cells recorded in MOs of naïve mice. **(d)** Median reaction times for different stimulus types, relative to mean over all stimulus types, for mouse behavior (grey, n=17, cf. Figure 1b), and the accumulator model fit to MOs activity in trained and naïve mice (solid and dashed black lines). **(e)** Mean behavior of the accumulator model fit to MOs neural activity in trained mice (large circles), together with mouse performance (small circles, n = 17, plotted as in Figure 1g). Solid lines represent the fit of the additive model to the accumulator model output. The accumulator model fits the mouse behavior significantly better than shuffled data (p < 0.01; shuffle test). **(f)** As in (c) but for an accumulator model trained on neural activity in MOs of naïve mice. There is no significant difference between the accumulator model and shuffled data (p > 0.05; shuffle test). **(g)** Simulation of right visual cortex inactivation, plotted as in (e). The accumulator model was applied after reducing the activity of visual-left-preferring cells reduced by 60% (large circles), plotted with the mean behavior from visual-cortex-inactivated mice (5 mice, small circles, cf. Figure 2g). The accumulator model fits the mouse behavior significantly better than shuffled data (p < 0.01; shuffle test). **(h)** Simulation of right MOs inactivation, plotted as in (e). Simply silencing all right hemisphere neurons did not provide a good fit to the observed behavior (Figure S7j), due to the mixture of left- and right-preferring neurons in each hemisphere. However, a model with connectional bias onto the downstream integrator, trained constraining neurons in the left and right hemispheres to have positive and negative weights, gave an accurate fit to MOs inactivation (5 mice, small circles, cf. Figure 2h) on reducing right-hemisphere activity by 60% (large circles). The accumulator model fits the mouse behavior significantly better than shuffled data (p < 0.01; shuffle test).

Although mouse behavior was not used to fit the model parameters, the model’s behavior matched that of the mice, but only when applied to recordings from trained mice. The model made a choice when *d*(*t*) crossed a decision boundary, and a weight vector ***w*** was found as that producing the fastest and most accurate choices possible given the MOs representation ***x***(*t*). The model reproduced the different behavioral reaction times for different stimulus types: faster in auditory and coherent trials, and slower in visual and conflict trials (Figure 6d; cf. Figure 1b). Furthermore, as observed with mice, the model integrated multisensory stimuli additively (Figure 6e; cf. Figure 1g). In contrast, an accumulator model trained on stimulus representations in naïve mice failed to reproduce mouse behavior, with no significant difference in model performance between shuffle and test data (p > 0.05; shuffle test Figure 6c,f). These results suggest that the spatial audiovisual signals we observed in MOs have the properties required to produce the behavior of the mice but appear in MOs only after the task has been learned.

The accumulator model even predicted the outcome of inactivation. Suppressing left-preferring visual neurons in our model reproduced the effects of inactivating right visual cortex (Figure 6g; cf. Figure 2g). However, a homologous suppression of all neurons recorded in the right hemisphere failed to reproduce the rightward bias observed in MOs inactivation (Figure S7j, cf. Figure 2h), because MOs neurons preferring either direction of stimulus are found equally in both hemispheres (Figure S7c-e). However, we could reproduce observed inactivation effects in a model where neurons projecting to the downstream integrator were those preferring contralateral stimuli (Li et al., 2015). We trained this model using the same method as before, but constraining weights from neurons in the left and right hemisphere to be positive and negative. This model predicted the lateralized effect of MOs inactivation (Figure 6h) on suppressing right-hemisphere activity. These results support the hypothesis that MOs neurons learn to additively integrate evidence from visual and auditory cortices, producing a population representation that is selectively sampled by a downstream circuit that makes decisions.

## Discussion

We found that in a task requiring multisensory integration of stimulus location, mice integrate auditory and visual cues additively, and that frontal area MOs is a key region for this additive integration. Inactivation of frontal cortex impaired audiovisual decisions, with strongest effects when the inactivation laser was targeted at anterior MOs. Recordings across frontal cortex found that representations of task variables were strongest in MOs. Sensory responses in MOs were present both inside and outside of the task context, but arose only after task training. MOs neurons combined visual and auditory location signals additively, and an accumulator model applied to MOs activity recorded in passive conditions in trained mice predicts the direction and timing of behavioral responses.

Taken together, our findings implicate MOs as a critical cortical region for integration of evidence from multiple modalities, at least in the current task. This is consistent with a general role for MOs in sensorimotor transformations: secondary motor cortex has been linked to multiple functions in rodents (Ebbesen et al., 2018), including flexible sensory-motor mapping (Barthas and Kwan, 2017; Duan et al., 2015), perceptual decision-making (Erlich et al., 2011, 2015; Guo et al., 2014; Hanks et al., 2015; Insanally et al., 2019; Zatka-Haas et al., 2021), value-based action selection (Sul et al., 2011), and the exploration-exploitation trade-off in both visual and auditory behaviors (Pisupati et al., 2021); furthermore, homologous regions of frontal cortex can encode multisensory information in primates (Gu et al., 2015). Sensory representations in rodent MOs have been seen to evolve with learning in visual tasks (Orsolic et al., 2019; Peters et al., 2021), consistent with our observations.

The effects of laser inactivation on responses to both modalities were strongest when the laser was aimed at anterior MOs. These inactivations may affect regions beyond those directly under the laser: a substantial degree of inactivation can be seen 1 mm from this location (Hao et al., 2021; Li et al., 2019; Zatka-Haas et al., 2021). Nevertheless, if the critical region for multisensory processing were some surface area other than anterior MOs, one would see a stronger effect targeting the laser there. It is also possible that targeting MOs inactivates regions below it, such as ACA or ORB. However, electrode recordings revealed that these other regions had no neural correlates of upcoming choice, and weaker correlates of stimulus location (excepting ACA where the difference did not reach significance).

We did observe an effect of inactivation targeted at the region between VISp and AUDp on both auditory and visual choices, and our current data cannot distinguish whether this reflects multisensory integration or simply lateral spread of the inactivation to both sensory cortices. The effects of inactivating this border region were weaker than those of inactivating MOs, particularly on auditory stimuli, as might be expected if the effect arose from diffusion of light through the brain to underlying auditory cortex. Furthermore, the effects of lateral cortex inactivation had a different character to those of MOs inactivation: MOs inactivation delayed choices on both sides, whereas inactivation of the border region had no significant effect on reaction times.

Our data cannot speak to a role of subcortical structures or non-frontal cortical areas below the surface (such as temporal association areas, ectorhinal area, or perirhinal area). However, we can conclude that the role of parietal cortex in this task is purely visual. This might appear to contradict previous work implicating parietal cortex in multisensory integration (Angelaki et al., 2009; Avillac et al., 2005, 2007; Fetsch et al., 2012; Gu et al., 2008; Khandhadia et al., 2021; Lippert et al., 2013; Nikbakht et al., 2018; Ohshiro et al., 2011; Raposo et al., 2014; Rohe and Noppeney, 2016; Seilheimer et al., 2013; Song et al., 2017), or showing multisensory activity in primary sensory cortices (Atilgan et al., 2018; Bizley and King, 2008, 2009; Driver and Noesselt, 2008; Ghazanfar and Schroeder, 2006; Ibrahim et al., 2016; Iurilli et al., 2012; Kayser et al., 2008; Meijer et al., 2017; Meredith and Allman, 2015). However, our finding agrees with evidence that parietal neurons can encode multisensory stimuli without being causally involved in a task (Cao et al., 2019; Gu et al., 2012; Licata et al., 2017; Raposo et al., 2014). We thus hypothesize that the causal role of visual and auditory cortices in this task is restricted to processing stimuli of the corresponding modality and relaying the signals to other regions (possibly via unimodal higher sensory areas (Oh et al., 2014)) where the two information streams are integrated (Cao et al., 2019; Jones and Powell, 1970; Rohe and Noppeney, 2016).

An additive integration strategy is optimal when the probability distributions of visual and auditory signals are conditionally independent given the stimulus location (Gold and Shadlen, 2002), but it may be a useful heuristic (Gardner, 2019) in a broader set of circumstances. In our task, independence in fact holds only approximately (Appendix 1). Nevertheless, additive integration is a simple computation (Ma et al., 2006), that does not require learning the detailed statistics of the sensory world, and gives performance close to the theoretical optimum under a wide range of situations. The additive model we observe in mice derives from Bayesian integration, the predominantly accepted integration strategy in humans and other animals (Alais and Burr, 2004; Angelaki et al., 2009; Chandrasekaran, 2017; Ernst and Banks, 2002; Fetsch et al., 2012; Franklin and Wolpert, 2011; Gu et al., 2008; Körding and Wolpert, 2004; Raposo et al., 2012; Sheppard et al., 2013; Witten and Knudsen, 2005). However, there is a useful distinction between the model used here and some previous work. In particular, many previous studies fit psychometric curves based on a cumulative Gaussian function (Ernst and Banks, 2002; Sheppard et al., 2013), which necessitates using lapse rates (Pisupati et al., 2021). Our approach instead starts with a conditional independence assumption, which implies that estimated probabilities are a logistic function applied to a sum of evidence from the two modalities (Appendix 1), but does not speak to the shape or linearity of these evidence functions. We found empirically that power functions of contrast approximate the data well, but this was not a necessary assumption (Appendix 1). Indeed, fitting a nonparametric variant of this additive model, which has a separate weight for each auditory location and visual contrast, captured subtle idiosyncratic differences in contrast tuning and slightly outperformed the parametric variant, albeit at the cost of more parameters.

Our finding of additive integration appears to contradict previous observations using an audiovisual detection task, which suggested that mice were auditory dominant (Song et al., 2017). Indeed, models allowing for auditory dominance did not improve over the additive model, even though they have more parameters. We hypothesize that this difference to previous work might arise from differences in the neural representation of stimulus onsets and locations. Our task required audiovisual localization, and we found that neural signals representing stimulus locations were combined additively in MOs, with differences in the encoding of auditory and visual stimulus location able to explain the mice’s earlier reactions to auditory stimuli. However, we also saw that neural signals encoding auditory onset were stronger and substantially earlier than neural signals encoding either visual onset or stimulus location from either modality. In a detection task (Song et al., 2017), these strong and early auditory onset signals might dominate behavior. In other words, mice might integrate audiovisual signals additively when tasked with localizing a source but be dominated by auditory cues when tasked with detecting the source’s presence.

The sensory code we observed in MOs has some apparently paradoxical features, but these would not prevent its efficient use by a downstream accumulator. First, a neuron’s preference for visual location showed no apparent relation to its preference for auditory location, consistent with reports from multisensory neural populations in primates (Gu et al., 2008) and rats (Raposo et al., 2014). Such “mixed selectivity” might result from random network connectivity, which can be computationally powerful, allowing downstream circuits to quickly learn to extract behaviorally relevant feature combinations (Caron et al., 2013; Fusi et al., 2016; Maass et al., 2002; Rigotti et al., 2013). Neurons encoding incoherent stimulus locations would not prevent a downstream decision circuit from learning to respond correctly; they could be ignored in the current task, but they would provide flexibility should task demands change. Second, although an approximately equal number of MOs neurons in each hemisphere preferred left and right stimuli of either modality, inactivation of MOs caused a lateralized effect on behavior. This apparent contradiction could be resolved if a specific subset of cortical neurons showed lateral bias (Li et al., 2015), or if the downstream decision circuit weighted MOs neurons in a biased manner. Indeed, midbrain neurons encoding choices in a similar task are highly lateralized (Steinmetz et al., 2019), and the subcortical circuits connecting MOs to midbrain stay largely within each hemisphere. Consistent with this hypothesis, if we constrained the accumulator model so that MOs neurons can only contribute to decisions toward the contralateral side, it reproduced the lateralized effects of MOs inactivation.

In summary, our data suggest that MOs neurons learn to additively integrate evidence from visual and auditory stimuli, producing a population representation that is suitable for guiding a downstream circuit that makes decisions by integration-to-bound. We hypothesize that this information is conveyed via sensory cortices to MOs, and is then fed to downstream circuits that accumulate and threshold activity to select an appropriate action (Figure 7). Based on results in a unisensory task (Steinmetz et al., 2019) we suspect these circuits may include neurons of basal ganglia and midbrain, and possibly recurrent loops including neurons within MOs itself.

**Figure 7.**
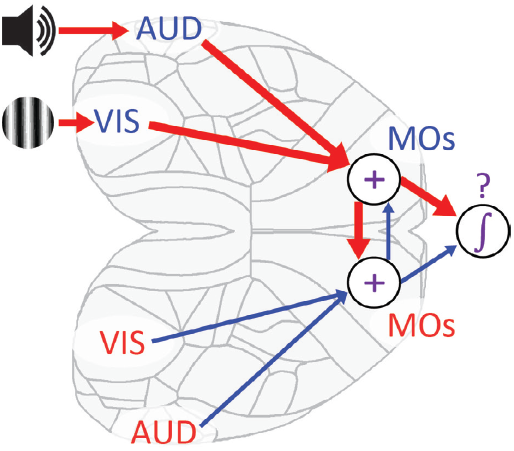
Diagram of hypothesized audiovisual integration pathway through cortex. Our data suggest that visual and auditory unisensory information are conveyed via sensory cortices (VIS and AUD) to MOs, where a bilateral representation results from interhemispheric connections. Downstream circuits accumulate MOs activity, with a biased sampling of neurons responding to contralateral stimuli, with an appropriate action determined by an integration to bound mechanism.

## Acknowledgements

We thank Michael Krumin, Peter Zatka-Haas, Andrew J Peters, Max Hunter, Flóra Takács, James Chadwick, Paul Johnson, and Ian Macartney for assistance with the experimental setup; Charu Reddy, Laura Funnell, Dylan Rich, Siddharth Kackar and Hamish Forrest for help with mouse husbandry, training, experimental assistance and spike sorting; Charu Reddy, Laura Funnell, and Flóra Takács for surgical assistance and optimization; Laura Funnell, Rakesh Raghupathy, David Orme, and Magdalena Robacha for histology processing; Samuel Picard, Célian Bimbard, and Maxwell Shinn for reading earlier versions of the manuscript. This work was supported by the Wellcome Trust (Sir Henry Wellcome Postdoctoral Fellowship 531073 to PC, and grants 205093 and 204915 to MC and KDH), the European Union’s Horizon 2020 research and innovation programme (European Research Council grant 694401 to KDH, and Marie Sklodowska-Curie fellowship 531949 to PC), and the Biotechnology and Biological Sciences Research Council (Responsive mode grant BB/T016639/1 to MC and PC). TS is supported by the Sainsbury Wellcome Centre PhD program. MC holds the GlaxoSmithKline/Fight for Sight Chair in Visual Neuroscience.

## Author contributions

PC, MC, and KDH conceived of and designed the study. PC designed the task. PC and MJW collected the behavioral data. PC collected the neural data. PC and TS analyzed data. PC, KDH, and MC wrote the manuscript with input from TS, and PC and KDH wrote the initial draft. Correspondence and material requests should be directed to Philip Coen.

|  | Coen | Sit | Wells | Carandini | Harris |
| --- | --- | --- | --- | --- | --- |
| Conceptualization | • |  |  | • | • |
| Methodology | • |  | • | • | • |
| Software | • | • | • |  |  |
| Formal Analysis | • | • |  |  | • |
| Investigation | • | • |  |  |  |
| Resources |  |  |  |  |  |
| Writing – Original Draft | • |  |  |  | • |
| Writing – Review & Editing | • |  |  | • | • |
| Visualization | • | • |  |  |  |
| Supervision | • |  |  | • | • |
| Funding Acquisition | • |  |  | • | • |

## Methods

Experimental procedures were conducted according to the UK Animals Scientific Procedures Act (1986) and under personal and project licenses released by the Home Office following appropriate ethics review.

### Terminology

Here, we define some terms used throughout the methods and manuscript. A “stimulus condition” refers to a particular combination of auditory and visual stimuli; for example, a visual stimulus of 40% contrast on the left and an auditory stimulus presented on the right. A “stimulus type” refers to a category that may comprise several stimulus conditions. We define five different stimulus types: unisensory auditory, unisensory visual, coherent, conflict, and neutral. “Unisensory auditory” trials are when an auditory stimulus is presented on the left or right, and visual contrast is zero (grey screen). “Unisensory visual” trials are when a stimulus of any contrast greater than zero is presented on the left or right, and the auditory stimulus is presented in the center (during behavior) or is absent (during passive conditions). “Coherent” trials are when a visual stimulus with non-zero contrast is presented on the same side as an auditory stimulus. “Conflict” trials are when a visual stimulus with non-zero contrast is presented on a different side from an auditory stimulus. “Neutral” trials are when the visual contrast is zero and the auditory stimulus is presented in the center. We refer to a single experimental recording (whether purely behavior, or combined with optogenetic inactivation or electrophysiology) as a “session.” Sessions can vary in duration and number of trials. Throughout the manuscript, “t-test” indicates a two-sided t-test unless otherwise specified. When referring to inactivation we use the term “site” to refer to a single target location (of which there were 52 in total) on dorsal cortex and “region” to refer to a collection of sites (3 sites in each case) in visual, lateral, somatosensory, or frontal cortex.

### Mice

Experiments were performed on 3 male and 15 female mice, aged between 9 and 21 weeks at time of surgery. For all experiments, we used either transgenic mice expressing ChR2 in Parvalbumin-positive inhibitory interneurons (Ai32 [Jax #012569, RRID:IMSR_JAX: 012569] x PV-Cre [Jax #008069, RRID:IMSR_JAX: 008069]) or wild type C57BL/6J [Jackson Labs, RRID:IMSR_JAX:000664]. 17 mice contribute to behavioral data (Figure 1), 5 mice contribute to optogenetic inactivation data (Figure 2), and 6/4 mice contribute to electrophysiological recordings in trained/naïve mice (Figure 3-6). Behavioral data (Figure 1) comprised both sessions without any optogenetic inactivation and non-inactivation trials within optogenetic experiments.

### Surgery

A brief (around 1 h) initial surgery was performed under isoflurane (1–3% in O2) anesthesia to implant a steel headplate (approximately 25 × 3 × 0.5 mm, 1 g) and, in most cases, a 3D-printed recording chamber. The chamber comprised two pieces of opaque polylactic acid which combined to expose an area approximately 4 mm anterior to 5 mm posterior to bregma, and 5 mm left to 5 mm right, narrowing near the eyes. The implantation method largely followed established methods (Guo et al., 2014) and has been previously described (Steinmetz et al., 2017). In brief, the dorsal surface of the skull was cleared of skin and periosteum. The lower part of the chamber was attached to the skull with cyanoacrylate (VetBond; World Precision Instruments) and the gaps between chamber skull were filled with L-type radiopaque polymer (Super-Bond C&B, Sun Medical). A thin layer of cyanoacrylate was applied to the skull inside the cone and allowed to dry. Thin layers of UV-curing optical glue (Norland Optical Adhesives #81, Norland Products) were applied inside the cone and cured until the exposed skull was covered. The head plate was attached to the skull over the interparietal bone with Super-Bond polymer. The upper part of the cone was then affixed to the headplate and lower cone with a further application of polymer. After recovery, mice were treated with carprofen for three days, then acclimated to handling and head-fixation before training.

### Audiovisual behavioral task

The two-alternative forced choice task design was an extension of a previously described visual task (Burgess et al., 2017). It was programmed in Signals, part of the Rigbox MATLAB package (Bhagat et al., 2020). Mice sat on a plastic apparatus with their forepaws on a rigid, rubber Lego wheel affixed to a rotary encoder (Kubler 05.2400.1122.0360). A plastic tube for delivery of water rewards was placed near the subject’s mouth.

Visual stimuli were presented using three computer screens (Adafruit, LP097QX1), arranged at right angles to cover ±135° azimuth and ±45° elevation, where 0° is directly in front of the subject. Each screen was roughly 11 cm from the mouse’s eyes at its nearest point and refreshed at 60 Hz. Intensity values were linearized (Burgess et al., 2017) with a photodiode (PDA25K2, Thor labs). The screens were fitted with Fresnel lenses (Wuxi Bohai Optics, BHPA220-2-5) to ameliorate reductions in luminance and contrast at larger viewing angles, and these lenses were coated with scattering window film (‘frostbite’, The Window Film Company) to reduce reflections. Visual stimuli were flashing vertical Gabors presented with a 9° Gaussian window, spatial frequency 1/15 cycles per degree, vertical position 0° (i.e. level with the mouse) and phase randomly selected on each trial. Stimuli flashed at a constant rate of 8Hz, with each presentation lasting for ~50 ms (with some jitter due to screen refresh times).

Auditory stimuli were presented using an array of 7 speakers (102-1299-ND, Digikey), arranged below the screens at 30° azimuthal intervals from −90° to +90° (where −90°/+90° is directly to the left/right of the subject). Speakers were driven with an internal sound card (STRIX SOAR, ASUS) and custom 7-channel amplifier (http://maxhunter.me/portfolio/7champ/). The frequency response of each speaker was individually estimated in situ with white noise playback recorded with a calibrated microphone (GRAS 40BF 1/4” Ext. Polarized Free-field Microphone). For each speaker, a compensating filter was generated to flatten the frequency response using the Signal Processing Toolbox in MATLAB. Throughout all sessions, we presented white noise at ~50 dbSPL to equalize background noise between different training and experimental rigs.

Auditory stimuli were 50 ms pulses of filtered pink noise (8-16kHz, 75-80 dbSPL), with 16ms sinusoidal onset/offset ramps. To ensure mice did not entrain to any residual difference in the frequency response of the speakers, auditory stimuli were further modulated on each trial by a filter selected randomly from 100 pre-generated options, which randomly amplified and suppressed different frequency components within the 8-16kHz range. As with visual stimuli, sound pulses were presented at a rate of 8Hz. On multisensory trials, the modulation of visual and auditory stimuli was synchronized, but software limitations and hardware jitter resulted in visual stimuli marginally preceding auditory stimuli by 10 *±* 12 ms (mean *±* s.d.).

A trial was initiated after the subject held the wheel still for a short quiescent period (duration uniformly distributed between 0.1 and 0.25 s on each trial; Figure 1a). Mice were randomly presented with different combinations of visual and auditory stimuli (Figure S1a). Visual stimuli varied in azimuthal position (−60° or +60°) and contrast (0%, 10%, 20%, 40%, and 80%, and also 6% in a subset of mice). On unisensory auditory trials, visual contrast was zero (grey screen). Auditory stimuli varied only in azimuthal position: −60°, 0°, or +60°; on unisensory visual trials, auditory stimuli were positioned at 0°. A small number of “neutral trials” had zero visual contrast, and an auditory stimulus at 0°. The ratio of unisensory visual/unisensory auditory/multisensory coherent/multisensory conflict/neutral trials varied between sessions but was ~ 10/10/5/5/1, and stimulus side was selected randomly on each trial. When a mouse was trained with 5 auditory azimuth locations (Figure S1k-l), the additional azimuths were −30° and +30°.

After stimulus onset there was a 0.5 s open-loop period, during which the subject could turn the wheel without penalty, but stimuli were locked in place and rewards could not be earned. The mice nevertheless typically responded during this open-loop period (Figure S1f). At the end of the open-loop period, an auditory Go cue was delivered through all speakers (10 kHz pure tone for 0.1 s) and a closed-loop period began in which the stimulus position (visual, auditory, or both) became coupled to movements of the wheel. Wheel turns in which the top surface of the wheel was moved to the subject’s right led to rightward movements of stimuli on the speaker array and/or screen, that is, a stimulus on the subject’s left moved towards the central screen. For visual or auditory stimuli, the position updated at the screen refresh rate (60Hz) or the rate of stimulus presentation (8Hz). In trials, where auditory stimuli were presented at 0°, the auditory stimulus did not move throughout the trial. A left or right turn was registered when the wheel was turned by an amount sufficient to move the stimulus by 60° in either azimuthal direction (~30° of wheel rotation, although this varied across mice/sessions); if this had not occurred within 1 s of the auditory Go cue, the trial was recorded as a “timeout.” On unisensory visual, unisensory auditory, and multisensory coherent trials, the subject was rewarded for moving the stimulus to the center. If these trials ended with an incorrect choice, or a timeout, then the same stimulus conditions were repeated up to a maximum of 9 times. In neutral and conflicting multisensory trials, left and right turns were rewarded with 50% probability (Figure S1a), and trials were only repeated in the event of a timeout, not an unrewarded choice. An incorrect choice or timeout resulted in an extra 2 s delay before the next trial for all stimulus conditions. After a trial finished (i.e. after either reward delivery or the end of the 2 s delay), an inter-trial interval of 1.5 to 2.5 s (uniform distribution) occurred before the software began to wait for the next quiescent period. Behavioral sessions were terminated at experimenter discretion once the mouse stopped performing the task (typically 1 h).

Mice were trained in stages (Figure S1b). First, they were trained to ~70% performance with only coherent trials; then auditory, visual, and neutral/conflict trials were progressively introduced based on experimenter discretion. Using this training protocol, ~80% of mice learnt the task, and those that did learn reached the final stage in < 30 sessions (Figure S1c).

### Behavioral quantification

With the exception of specific analyses of timeout trials (Figure 2l, Figure S5i-p), timeouts and repeats following incorrect choices were excluded. To remove extended periods of mouse inattention at the start and end of experimental sessions, we excluded trials before/after the first/last three consecutive choices without a timeout. The 6% contrast level was included in analyses of inactivation experiments (Figure 2) as all mice contributing to these analyses were presented with 6% contrast levels, but not all behavioral and electrophysiology sessions included 6% contrast.

On 91.5% of trials (142853/156118), subjects responded to the stimulus onset by turning the wheel within the 500 ms open-loop period (Figure S1f). For data analysis purposes, we therefore calculated mouse choice and reaction time from any wheel movements after stimulus onset (Figure S1d-e), even though during the task, rewards would only be delivered after the open-loop period had ended. These choices were defined by the first time point at which the movement exceeded ~30° of wheel rotation (the exact number varied across sessions/mice, Figure S1d), the same threshold required for reward delivery during the closed-loop period. This matched the outcome calculated during the closed loop period on 94.9% of trials (148203/156118). The reaction time was defined as the last time prior to the choice threshold at which velocity crossed 0 after at least 50 ms at 0 or opposite to the choice direction, and then exceeded 20% of the choice threshold per second for at least 50 ms (Figure S1e). On 5.1% of trials (8380/164498), no such timepoint existed or movement was non-zero within 10 ms of stimulus onset; these trials were excluded. On 38% of trials (59498/156118), mice made sub-threshold movements prior to their calculated reaction time. To eliminate the possibility that these earlier movements were responsible for the neural decoding of choice (Figure 3g) we repeated this analysis using only trials without any movement prior to the calculated reaction time (Figure S6c), which did not change the results.

When calculating performance for each stimulus type for a single visual contrast (Figure 1e, Figure S1n,p), the value for each mouse was calculated within each session before taking the mean across sessions. We then took the mean across symmetric presentations of each stimulus condition (e.g. unisensory auditory left and right trials). In the case of reaction time (Figure S1h-j), we calculated the median for each session before taking the mean across sessions and symmetric presentations. For relative reaction time (Figure 1b, Figure S1o,q, Figure 6d) we also subtracted the mean across all stimulus types for each mouse. For both performance and reaction time, differences between stimulus types were quantified with a paired t-test (n = 17 mice). Using this analysis, we established that reaction times were faster on unisensory auditory trials than unisensory visual trials (Figure 1b). To confirm that the earlier movements on unisensory auditory trials were genuine choices rather than reflexive movements unrelated to the stimulus location, we predicted whether stimuli were presented on the right or left in unisensory auditory and unisensory visual trials from the wheel velocity at each timepoint after stimulus onset. Trial data was subsampled for each session (to equalize the number of stimuli appearing on the left and right) and split into test and training data (2-fold cross validation). Mean prediction accuracy was calculated by first taking the mean across sessions, then across mice. Consistent with our conclusions from calculated reaction times, auditory location could be decoded earlier than visual location (Figure S1g). This conclusively demonstrates that mice were able to identify the location of an auditory stimulus earlier than a visual stimulus.

### Video motion energy analysis

Because neural activity across the brain is related to bodily motion (Musall et al., 2019; Stringer et al., 2019), we asked if mice still respond to stimuli in the passive condition. We filmed the mouse at 30 frames per second (DMK 23U618, The Imaging Source). We quantified the motion energy on each trial by averaging the absolute temporal difference in the pixel intensity values, across all pixels in a region of interest including the face and paws, and across a time period 0 to 400 ms after stimulus onset, which typically included the mouse response during behavior (Figure S1f). This analysis established that mice exhibit minimal movement in response to task stimuli during passive conditions (Figure S7h).

### Optogenetic inactivation

For optogenetic inactivation experiments (Figure 2, Figure S4-5) we inactivated several cortical sites through the skull using a blue laser (Cardin et al., 2009; Guo et al., 2014; Olsen et al., 2012; Zatka-Haas et al., 2021), in transgenic mice expressing ChR2 in Parvalbumin-expressing inhibitory interneurons (Ai32 x PV-Cre). Unilateral inactivation was achieved using a pair of mirrors mounted on galvo motors (GVSM002-EC/M, Thor labs) to orient the laser (L462P1400MM, Thor labs) to different points on the skull. On every trial, custom code drove the galvo motors to target one of 52 different coordinates distributed across the cortex (Figure 2a), along with 2 control targets outside of the brain (Figure S4c). A 3D-printed isolation cone prevented laser light from reaching the screens and influencing behavior. Inactivation coordinates were defined stereotaxically from bregma and were calibrated on each session. Anterior-posterior (AP) positions were distributed across 0, ±1, ±2, ±3, and −4 mm. Medial-lateral (ML) positions were distributed across ±0.6, ±1.8, ±3.0, and ±4.2 mm. On 75% of randomly interleaved trials, the laser (40 Hz sine wave, 462 nm, 3 mW) illuminated a pseudorandom location from stimulus onset until the end of the response window 1.5 s later (both open and closed loop periods, irrespective of mouse reaction time). The laser was not used on trial repetitions due to incorrect choices or timeouts. Pseudorandom illumination meant that a single cortical site was inactivated on only 1.4% of trials per session. This discouraged adaptation effects but required combining data across sessions for analyses. The galvo-mirrors were repositioned on every trial, irrespective of whether the laser was used, so auditory noise from the galvos did not predict inactivation.

To investigate the effects of inactivation at different time points (Figure 2j-k), in separate experiments the laser was switched on for 25 ms (DC) at random times relative to stimulus onset (−125 to +175 ms drawn from a uniform distribution). Inactivation was randomly targeted to visual areas (VISp; −4 mm AP, ±2 mm ML) or secondary motor area (MOs; +2 mm AP, ±0.5 mm ML) on 25% of trials.

### Psychometric modelling

The model we use throughout the text, the (parametric) additive model, is given by the equation

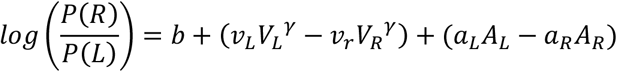

Model parameters were fit by maximizing the likelihood of observed behavioral data using MATLAB’s *fmincon* function to implement the *interior-point* algorithm to find 6 fit parameters: *v_R_, v_L_ a_R_* and *a_L_* representing sensitivities to right and left visual and auditory stimuli, *b* representing bias, and the contrast gain parameter *γ*. When fitting for individual mice (Figure 1c,f, Figure S1k, Figure S3a-o), models were fit to data combined across sessions. When the model was fit to combined data from multiple mice (Figure 1d,g, Figure 2g-h, Figure S2, Figure S4e-h), trials were subsampled to equalize numbers across mice before fitting the model. This subsampling process was repeated 10 times, and plots reflect the mean model parameters, and fraction of rightward choices, across repeats. For visualization, if the log-odds were not defined for a given stimulus condition (because a mouse, or mice, made only rightward or leftward choices) the log odds were regularized by adding one trial in each direction. This was only necessary for the coherent stimulus condition at 10% contrast in Figure 2h.

We compared our additive model to a range of other models (Figure S2), all fit the same way. The “auditory-only” model (Figure S2a) was given by

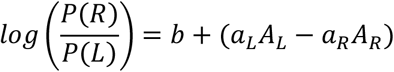

And the “visual-only” model (Figure S2b) was given by:

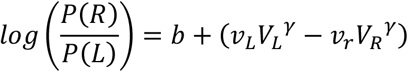

For the “auditory dominance” model (Figure S2c), we set the visual weight to zero whenever auditory and visual stimuli were in conflict:

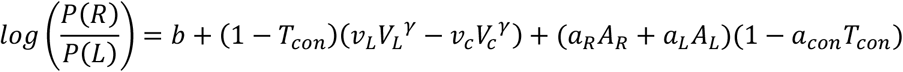

Here, *T_con_* is a binary variable, equal to 1 or 0 to indicate whether each trial is a multisensory conflict trial, and *a_con_* is an additional fit parameter. We tested this model both with *a_con_* = 0 (Figure S2c) and with *a_con_* allowed to take any value (Figure S2d).

As a more general test for any evidence of visual or auditory dominance during audiovisual trials, we fit a “sensory bias” model (Figure S2e) with additional auditory and visual weights on coherent and conflict trials:

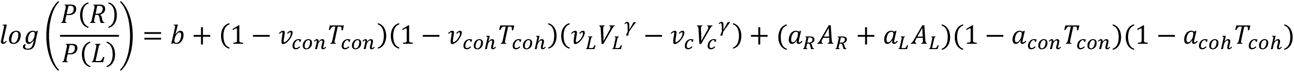

Here, *a_con_*, *v_con_, a_coh_*, and *v_coh_* are fit parameters and *T_coh_* is a binary variable, equal to 1 or 0 to indicate whether each trial is a multisensory coherent trial. Our 6-parameter additive model is a special case of this 10-parameter sensory bias model, when the 4 multisensory parameters are zero.

The 11-parameter “additive nonparametric” model (Figure S2f) is similar to the usual additive model, but can fit any function of visual contrast, not just a power function:

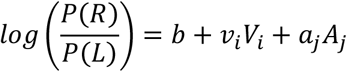

Here *V_i_* and *A_j_* are binary variables indicating the presence of visual contrast *i* and auditory location *j* on a given trial. The parameters *V_i_* and *a_j_* represent the visual and auditory sensitivities to visual contrast *i* and auditory location *j*, and are constrained to be 0 for zero contrast visual and central auditory stimuli.

Finally, to determine whether a generic non-additive model of multisensory integration could improve model fit, we tested a 27-parameter “full model” which had a weight for each combination of auditory and visual stimulus (Figure 1h, Figure S3p).

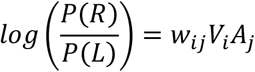

We evaluated the fit of each model by its log_2_-likelihood ratio relative to a bias-only model *log*(*p*(*R* | *A,V*)/*p*(*L* | *A, V*)) = *b* using 5-fold cross-validation. After normalizing by the number of trials, this yields a quantity in bits per trial: the number of bits two parties would save in communicating the mouse’s choice, if the stimulus is known to both. We compared all models to the additive parametric model (Figure 1h, Figure S2). Across 17 mice, the additive model was not significantly worse than the full model, either when trained on all trial types (Figure 1h), or when trained only on unisensory and neutral trials but tested on all trials including multisensory combinations (Figure S3p).

When fitting the additive model to data where different regions of dorsal cortex were inactivated, three target locations were combined to represent each region. For visual cortex (Figure 2g,i,l, Figure S4h, Figure S5), these were (−4,1.2), (−4,3) and (−3,3), where coordinates indicate (AP, ML) distances from bregma in mm. For frontal cortex (2,0.6), (2,1.8) and (3,0.6) (Figure 2h,i,l, Figure S4g-h, Figure S5); for lateral areas proximal to auditory cortex (−4,4.2), (−2,4.2), (−2,4.2) (Figure S4e,h, Figure S5); for areas proximal to somatosensory cortex (1,3), (0,3), (0,4.2) (Figure S4f,h). When fitting these models, the contrast gain parameter was fixed at the value obtained when fitting to non-inactivation trials.

For a mouse presented with 5 auditory conditions (Figure S1k-l), the additive model contained two additional auditory parameters, such that each non-zero auditory azimuth had a distinct weight:

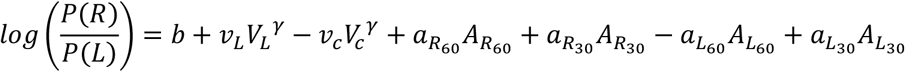

Here, *R*_60_, *R*_30_, *L*_60_, and *L*_30_ indicate whether the auditory stimulus was presented at 30° or 60° on the left or right.

### Quantifying effects of optogenetic inactivation on choice

To quantify the change in the fraction of rightward choices when a particular cortical location was inactivated, we used a shuffle test (Figure 2b-e, Figure S4a-b). Data were initially combined across 5 mice and segregated by stimulus type (unisensory visual, unisensory auditory, multisensory coherent, or multisensory conflict). For each type, data were further segregated into non-inactivation trials (laser off) and inactivation trials (laser on) grouped by the targeted area of dorsal cortex. For trials where the stimulus was presented on the right, we reversed the laterality of the stimulus and inactivation location such that all stimuli were effectively presented on the left (visual stimulus in the case of conflict trials). Data were randomly subsampled to equalize the number of trials contributed by each mouse to non-inactivation and the inactivation trials at each targeted location. We then calculated the difference in the fraction of rightward choices for each targeted location compared with non-inactivation trials. This process was repeated 25,000 times with different subsampling to produce a mean change in fraction of rightward choices for each inactivated location on dorsal cortex.

For each of the 25,000 iterations, we proceeded to generate 10 independent shuffles, where the labels for targeted location and trial identity (inactivation or non-inactivation) where randomly reassigned. We thus generated a null distribution for each targeted location, comprising 250,000 datapoints from independent shuffles. For each targeted location, the position of the unshuffled result within this null distribution gave the significance value for that location (e.g. top/bottom 0.05% for *p* < 0.001, top/bottom 0.005% for *p* < 0.0001).

When assessing the symmetry of inactivation effects across hemispheres (Figure S4a) the process was as described above, but without reversing the laterality of any trials. To confirm results were similar across mice (Figure S4b), we repeated this process for individual mice. In this case, the number of shuffled iterations remained at 250,000 but no subsampling was required (because there was no need to equalize across mice).

To test how pulsed inactivation at different times affected choices (Figure 2j-k), data were combined across 7 mice. Trials where stimuli appeared on the right were reversed such that an increase in the fraction of rightward choices corresponded to an increase in the fraction of ipsilateral choices. Experimental sessions with fewer than 75 inactivation trials were excluded to ensure that each session contributed to both the fraction of inactivation and control trials. Laser onsets were binned using a sliding 70 ms boxcar window, and the time between stimulus onset and inactivation was defined as the center of this window. In each 70 ms time window, we calculated the change in fraction of rightward choices compared with non-inactivation trials, and the significance of this difference was established with a Fisher’s exact test. Each timepoint was defined as significant if it, or both its neighboring timepoints, passed the significance criterion of p < 0.001.

### Quantifying effects of optogenetic inactivation on model parameters

To quantify the changes in parameters of the additive model (Figure 2f, Figure S4d) the analysis closely mirrored the steps described above, but trial types were not segregated by stimulus type. The additive model was reparametrized such that stimuli were defined as being ipsilateral or contralateral to the site of inactivation, effectively combining data across hemispheres:

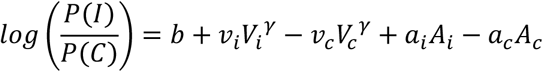

Here, *V_c_* and *V_i_* are contralateral and ipsilateral visual contrasts, and *A_c_* and *Ai* are contralateral and ipsilateral auditory azimuths. *v_i_*, *v_c_ a_i_* and *a_c_* represent sensitivities to contralateral and ipsilateral visual and auditory stimuli, while *b* represents the bias, and *γ* the contrast gain parameter. The unshuffled dataset comprised 2,500 different subsamples, and in each iteration, we fit the additive model to the non-inactivation data and to the inactivation data for each targeted location. This gave the mean change in each model parameter at each location on dorsal cortex. We compared this value to a null distribution (generated as described above, total of 25,000 independent shuffles) to establish the significance of each change. Since we observed no change in the contrast gain parameter, *γ* (Figure S4d), in our final analysis we fixed this value according to the non-inactivation trials and only quantified changes in the remaining 5 parameters (Figure 2f).

To determine whether inactivating these regions caused a significant change in model parameters compared with non-inactivation trials, we evaluated the loglikelihood ratio between a model trained and evaluated on inactivation trials and a model trained on non-inactivation trials and then evaluated to inactivation trials for individual mice. We then determined if the loglikelihood ratio was significantly different from zero across the 5 mice using a t-test (Figure 2g-h, Figure S4e-f).

To test whether inactivation of the four different regions (frontal, visual, lateral sensory, and somatosensory cortices) had significantly different effects we used a shuffle test to evaluate data combined across all mice (Figure S4g). For each pair of regions, as well as non-inactivation trials, we calculated inter-region loglikelihood (where the model was fit to trials from one region and then evaluated on another region) and a within-region loglikelihood (where the model was trained and evaluated on data from one inactivated region). We repeated this process in 100 different subsamples, equalizing the number of trials from each mouse, and the number of trials in the train and test sets, and took the mean loglikelihood ratio between the inter-region and within-region results. We then generated a null distribution by repeating this process, but with the label of the inactivation site shuffled before splitting the data to perform the inter-region and within-region comparison (total of 1,000 independent shuffles). For each pairwise regional comparison, we compared the mean unshuffled loglikelihood ratio to the null distribution and found that every inter-region loglikelihood was significantly lower than the within-region loglikelihood (p < 0.05, Bonferroni-corrected).

### The effect of inactivation on reaction time and fraction of timeout trials

We used a linear mixed effects model (LME) to determine the effect of inactivating visual, lateral, or frontal cortices on mouse reaction time for each stimulus type (auditory, visual, coherent and conflicting) when stimuli were contralateral or ipsilateral to the site of inactivation (Figure 2i, Figure S5a-h). For each mouse, we computed the median reaction time over trials of all sessions for each combination of stimulus condition and inactivation region. We fit the following LME model to this data using MATLAB’s *fitlme* function:

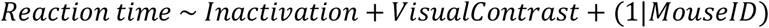

Here, *Reaction time* is the response variable, *Inactivation* (binary) and *VisualContrast* (categorical) were fixed effect terms, and *MouseID* was a random effect on the intercept. We separate LMEs for each stimulus type and region of inactivation. In each case, we assessed the sign and significance of the *Inactivation* term to assess the impact of inactivation on mouse reaction time (Figure 2i, Figure S5a-h). To make direct inter-region comparisons, we modified the LME model:

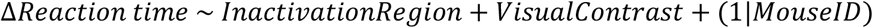

Here, *ΔReaction time* is the difference in reaction time between the inactivated trials and non-inactivation trials for each stimulus condition (for each stimulus condition within a stimulus type). *InactivationRegion* is a binary fixed effect term which identifies which of the two inactivation regions being compared (for example, frontal and visual, Figure 2i) was targeted. As above, we assessed the sign and significance of the *InactivationRegion* term to determine whether the inactivation region had a significant effect on the change in reaction time (Figure 2i, Figure S5a-h).

Statistical analyses of timeout trials were performed in the same way as the two previous LMEs, but *Reaction time* was replaced with *Timeout* (a binary response variable representing whether the mouse gave a response within 1.5 s) and *ΔReaction time* was replaced with *ΔTimeouts*, a continuous response variable representing the difference in timeout frequency between the inactivated trials and non-inactivation trials (Figure 2l, Figure S5i-p).

### Neuropixels recordings

Recordings in behaving mice were made using Neuropixels (Phase3A; (Jun et al., 2017)) electrode arrays, which have 384 selectable recording sites out of 960 sites on a 1 cm shank. Probes were mounted to a custom holder (3D-printed polylactic acid piece) affixed to a steel rod held by a micromanipulator (uMP-4, Sensapex Inc.). Probes had a soldered external reference connected to ground which was subsequently connected to an Ag/AgCl wire positioned on the skull. On the first day of recording mice were briefly anaesthetized with isoflurane while one or two craniotomies were made with a biopsy punch. After at least three hours of recovery, mice were head-fixed in the usual position. The craniotomies, as well as the ground wire, were covered with a saline bath. One or two probes were advanced through the dura, then lowered to their final position at approximately 10 μm/s.

Electrophysiological data were recorded with Open Ephys (Siegle et al., 2017). Raw data within the action potential band (1-pole high-pass filtered over 300 Hz) was denoised by common mode rejection (that is, subtracting the median across all channels), and spike-sorted using Kilosort 2 (www.github.com/MouseLand/Kilosort2). Units were manually curated using Phy to remove noise and multi-unit activity (Rossant et al., 2016). Each cluster of events (‘unit’) detected by a particular template was inspected, and if the spikes assigned to the unit resembled noise (zero or near-zero amplitude; non-physiological waveform shape or pattern of activity across channels), the unit was discarded. Units containing low-amplitude spikes, spikes with inconsistent waveform shapes, and/or refractory period contamination were labelled as ‘multi-unit activity’ and not included for further analysis.

To localize probe tracks histologically, probes were repeatedly dipped into a centrifuge tube containing DiI before insertion (ThermoFisher Vybrant V22888 or V22885). When probes were inserted along the same trajectory for multiple sessions (Figure S6a), they were coated with Dil on the first day, and subsequent recordings were estimated to have the same trajectory within the brain (although depth was independently estimated, Figure S6b). After experiments were concluded, mice were perfused with 4% paraformaldehyde. The brain was extracted and fixed for 24 h at 4 °C in paraformaldehyde before being transferred to 30% sucrose in PBS at 4 °C. The brain was then mounted on a microtome in dry ice and sectioned at 80 μm slice thickness. Sections were washed in PBS, mounted on glass adhesion slides, and stained with DAPI (Vector Laboratories, H-1500). Images were taken at 4× magnification for each section using a Zeiss AxioScan, in two colors: blue for DAPI and red for DiI. Probe trajectories were reconstructed from slice images (Figure S6a) using publicly available custom code (http://github.com/petersaj/AP_histology (Peters et al., 2021)). For each penetration, the point along the probe where it entered the brain was manually estimated using changes in the LFP signal (Figure S6b). Recordings were made in both left (47 penetrations) and right (41 penetrations) hemispheres. The position of each recorded unit within the brain was estimated from its depth along the probe. For visualization, the recorded cells were mapped onto a flattened cortex using custom code (Figure 3a). Given the small size the frontal pole (FRP), neurons in this region could not be confidently separated from MOs, and so were considered part of MOs for the purpose of this manuscript (14% of MOs cells; excluding these cells did not significantly impact results).

For recordings from naïve mice (Figure 5, Figure 6c-d,f), data were acquired with 4-shank Neuropixels 2.0 probes, which have 384 selectable recording sites out of 5000 sites on 4 1 cm shanks (Steinmetz et al., 2021). We recorded from the 96 sites closest to the tip of each shank. Electrophysiological data for these experiments were recorded with SpikeGLX (https://billkarsh.github.io/SpikeGLX/). The same procedures were followed as above for mouse surgery and manual curation of units. Changes in LFP signal were used to detect the point at which the probe entered the brain, and only cells within 1.25mm of the brain surface (i.e. within MOs) were included in analyses.

### Passive stimulus presentation recordings

Mice were presented with task stimuli under passive conditions after each behavioral recording session. Stimuli were presented in open-loop (entirely uncoupled from wheel movement) and mice did not receive rewards. Unisensory auditory, unisensory visual, coherent, and conflict trials were presented to mice. However, on unisensory visual trials, the auditory amplitude was set to zero (rather than positioned at 0° as in the task) to ensure visual sensory responses could be isolated. Due to time constraints, only one coherent and conflicting stimulus combination were presented (80% visual contrast in both cases), and the trial interval was reduced (randomly selected from 0.5 to 1 s). Stimulus conditions were randomly interleaved, and each condition was repeated ~50 times.

### Estimating firing rate

Unless otherwise specified, firing rates were calculated on each trial by binning in 2 ms windows and smoothing with a half-Gaussian filter with standard deviation of 60 ms. PSTHs were calculated by averaging this rate across trials.

### Decoding stimuli and choices from population activity

To decode stimuli and choices from neural activity (Figure 3e-g, Figure S6c-d), we trained a linear SVM decoder on the firing rate vector time-averaged over a window 0-300 ms after stimulus onset (Figure 3e-f), 0-130 ms before movement onset (Figure 3g, Figure S6c), or 150-300 ms after movement onset (Figure S6d). SVMs were trained separately for each Neuropixels behavioral recording, for any brain region with a minimum of 30 neurons recorded in that session. If more than 30 neurons were recorded, we repeatedly (5 repeats) selected a 30-neuron subset for decoding analysis and took the mean accuracy (5-fold cross-validated) across these repeats. Sessions with fewer than 25 trials of each decoded condition (e.g. left and right stimulus locations) were excluded. In the case of decoding visual location (Figure 3e), only trials with high-contrast (40% and 80%) stimuli were included. In each session, decoding accuracy was quantified as the fraction of test-set trials classified correctly, relative to the same number for a model with no access to the spike trains (whose optimal behavior is to always predict the most common stimulus on the training set):

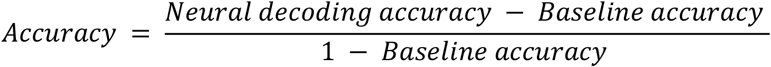

To compare the decoding accuracy between brain regions, we first performed a one-way ANOVA, which showed a significance difference (visual location: F=26.1, p < 10^-20^, auditory location: F=77.7, p < 10^-67^, and upcoming choice: F=21.0, p < 10^-13^). To compare pairwise differences, we fitted a linear mixed effects model:

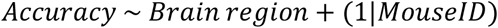

Here, *Accuracy* is defined as above, *Brain region* is a categorical fixed effect and *MouseID* was a random effect on the intercept, to takes account of the potential confound of differences in decoding accuracies across mice (Figure 3e-g).

### Combined-conditions choice/stimulus probability analysis

To quantify the selectivity of a cell for a choice while controlling for effects of stimulus (Figure S6h), we used the combined-conditions choice probability (ccCP, (Steinmetz et al., 2019)). This is based on an extension of the Mann-Whitney U statistic, defined as the fraction of pairs of trials of identical stimulus conditions but different choices, for which the firing rate on the right choice trial exceeds the firing rate on the left choice trial. The significance of this test statistic was evaluated by shuffling using a p-value of 0.01, meaning that the observed value has to be either below the 0.5 percentile or above the 99.5 percentile of a null distribution generated from 1000 shuffles of the choice labels for each stimulus condition in order to be deemed significant. For ccCP, we compared the firing rate averaged over 0 – 130 ms before movement onset between trials where the mouse made a leftward or rightward choice trials (Figure S6h).

To test for selectivity to one stimulus while controlling for the other stimulus and choice (Figure S6g), we used an analogous method, referred to as the combined conditions stimulus probability (ccSP). For visual ccSP, we compared the firing rate time-averaged over a 0 – 300 ms window after stimulus onset, between trials where the visual stimulus was on the left and trials where the visual stimulus was on the right, including only trials with high (40% or 80%) contrast (Figure S6g, left). For auditory ccSP, we compared the firing rate averaged over a time window 0 – 300 ms after stimulus onset between auditory-left and auditory-right trials (Figure S6g, right).

### Modelling neural activity

To predict firing rate time courses from task events (Figure 4a-c), we used an ANOVA-style decomposition. For this analysis, we pooled multisensory coherent and conflict trials of visual contrast 40% and 80% (using a single visual contrast did not impact results), resulting in four possible stimulus conditions: one for each combination of auditory and visual location. We defined binary variables *a_i_, v_i_, c_i_* = ±1 encoding whether auditory stimuli, visual stimuli, and choices are to the left or right on trial *i*. We can decompose ***F**_i_*(*t*), the firing rate vector on trial *i* at time *t* after stimulus onset, as:

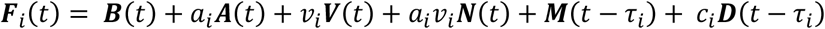

This model decomposes the response into a sum of 6 temporal kernels. ***B*** represents the grand mean stimulus response; ***A*** and ***V*** represent the additive main effects of auditory and visual stimulus location, and ***N*** represents a non-additive interaction between them. To account for the effects of movement, ***M*** is a kernel representing the mean effect of movement (relative to *τ_i_*, the time of movement onset on trial *i*) and ***D*** represents the effect of movement direction. ***B, A, V, N*** were allowed to be non-zero for −50 ≤ *t* ≤ 400 ms. ***M, D*** can be non-zero for −200 ≤ *t −τ_i_* ≤ 700 ms. Only trials with *τ_i_* < 300 ms were included. The model was fit using ridge regression with a regularization strength of *α=* 10, which we found to give optimal prediction accuracy. We fit this model to each neuron in MOs with a non-zero firing rate during behavior (n = 2183 neurons), using a training set consisting of half the trials (randomly selected). The error, *E*, of this fit was measured as:

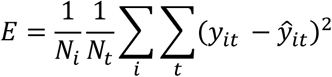

Here, *ŷ_it_* and *y_it_* are model prediction and test-set recorded firing rate on trial *i* and timepoint *t*, *N_i_* is the number of neurons, and *N_t_* is the number of time bins, spanning 0 to 400 ms relative to stimulus onset. *E* is thus the cross-validated mean-squared error between the predicted and the actual smoothed firing rate over this time window. To test for an additive code, we then repeated this process for an additive neural model where ***N*** = 0 (Figure 4c).

To investigate whether there was an interaction between stimulus condition and choice-related response, we also fit a model with 8 movement-aligned kernels, i.e. a movement and a direction kernel for each combination of the four possible audiovisual stimuli:

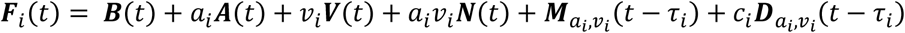

We compared this full model to the additive neural model (two movement kernels and ***N*** = 0) using the method described above (Figure S7a).

To model neural activity during passive stimulus presentation (Figure 4d-g), we used a reduced model without movement-aligned kernels:

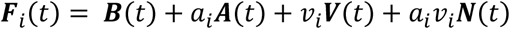

Here, only multisensory coherent and conflict trials of a single (80%) visual contrast were included (due to time constraints, this was the only contrast presented on multisensory trials in passive conditions). To test for an additive code, we repeated the process described above (on 2509 cells with non-zero firing rates (Figure 4f). No regularization was used for this analysis of passive data as it did not improve fits.

To compare the fit of linear and non-linear models of neural firing (Figure 4c,f, Figure S7a), we used a linear mixed effects method to determine the main effects of the prediction model, accounting for systematic differences in model fit across mice and across experiments within each mouse. This was done using the *fitlme* in MATLAB with the following formula:

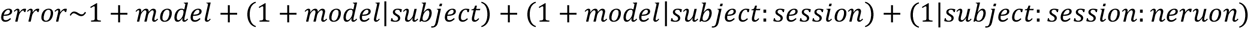

The error term *E* is modelled with an intercept, a fixed effect of the model type being used (e.g. either the additive or full model), random effects for the intercept and model type grouped by subjects, random effects for the intercept and model type grouped by session nested within subjects, and random effects for the intercept grouped by neurons nested within sessions within subjects. For all statistical tests we report the p-value of the main effect of the model type on the observed error values.

To examine the distribution of auditory and visual spatial sensitivity across neurons we used neural recordings from passive stimulus presentation (Figure 4g, Figure S7b). We selected neurons where the additive neural model (***N*** = 0) explained a minimum of 2% variance. For each neuron, we averaged the amplitude of the ***A*** and ***V*** kernels over a time window from 0 to 300 ms after stimulus onset (the kernels were fit using all trials). To test for a significant correlation between the signed magnitude of these time-averaged ***A*** and ***V*** kernels, we used the linear mixed effects model described above, but with *time-averaged **V**kernel* and *time-averaged A kernel* substituted for *error* and *model* (Figure 4g). To test for a relationship between the absolute values of the two kernels, we repeated this procedure but using the absolute, rather than the signed, time-averaged kernels (Figure S7b).

### Lateralization of stimulus and movement activity

To investigate whether there is lateralization in the spatial preference of auditory neurons, we examined time-averaged value of the *A* kernels (0 ms to 300 ms after stimulus onset) after fitting the additive model (*N* = 0) under passive conditions. We selected neurons for which the additive model performed better than a model with visual kernel alone, and compared the mean value of the *A* kernel for neurons recorded in each hemisphere (Figure S7d). We repeat the same procedure for the visual kernel weights to examine lateralization of visual spatial preference (Figure S7c).

To investigate the lateralization of movement-related responses, we repeated this procedure, but for the additive model (*N* = 0) during behavior. We then included all for which the directional movement kernel, *D* improved cross-validated fits. Mean kernel values of selected neurons were calculated using a time window −200 to 400 ms relative to movement onset (Figure S7e).

Statistical analysis to determine the lateralization of sensory and movement responses were performed with linear mixed effects model as described above, but with *time-averaged kernel* and *hemisphere* substituted for *error* and *model*.

### Quantifying single-neuron discrimination time

To identify when visual and auditory information began to be encoded in MOs (Figure 4h), we analyzed responses to passive unisensory stimuli. We first used permutation tests to select neurons sensitive to the presence (On-Off) and/or the location (Right-Left) of auditory and visual stimuli. To identify On-Off neurons, we calculated two PSTHs, one for sounds in each location, in a window 0 to 300 ms after stimulus onset, and computed the difference between the maximum of this PSTH and the mean firing rate 300 to 0 ms before stimulus onset. We compared this value to a null distribution obtained from 1000 shuffles of the pre/post-stimulus windows independently for each trial. A neuron was defined as significantly responding to a stimulus if the maximum difference in unshuffled data was in the 1st or 99th percentile of the null distribution for either left or right stimuli. For Right-Left neurons, the same method was uses, but using the maximum difference between the PSTHs for left and right auditory or visual presentations 0 to 300 ms after stimulus onset, and shuffling the left/right trial labels. This method identified 72 auditory (3%) and 68 visual (3%) Right-Left neurons.

For identified On-Off neurons, we calculated the discrimination time by separately comparing the pre- and post-stimulus firing rate in a sliding window of 50 ms with step size 5 ms, defining significance using a Mann-Whitney U test at p < 0.01, and requiring three consecutive significant time windows to qualify as the discrimination time. We excluded discrimination times that occurred more than 300 ms after stimulus onset as they are unlikely to be stimulus-related activity. This analysis was done separately for left and right stimuli, taking the earliest statistically significant time window in either stimulus condition. For identified spatially selective (Right-Left) neurons, we defined the discrimination time as the earliest time after stimulus onset where there is a significant difference in the response to left and right stimuli (Figure 4h). This method identified discrimination times for 82 and 36 auditory and visual On-Off neurons, and 59 and 36 auditory and visual Right-Left neurons. For each neuron we also calculated the 5-fold cross-validated decoding accuracy, relative to a baseline model (which always predicts the most-frequent stimulus-condition in the training set, as in Figure 3e-f), from the time-averaged firing rate in a window 0 to 100ms after the discrimination time using a linear SVM decoder (Figure S7i).

### Quantifying single-neuron discriminability index

To quantify single-neuron selectivity for sensory location and upcoming choice, we calculated the discriminability index (*d’* or d-prime) between different trial conditions (Figure 5, Figure S6e-f). The discriminability index is defined as:

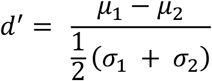

Here *μ_1_* and *μ_2_* are the mean firing rate of the neuron over a stimulus onset for quantifying stimulus responses, or (0 – 130 ms before movement onset for quantifying choice coding), and *σ*_1_ and *σ*_2_ are the standard deviation of the firing rate across the respective trial conditions.

To compare single neuron discriminability indices across brain regions, we first performed a one-way ANOVA on the mean of the absolute value of the discriminability index across neurons of each recorded session for each brain region, which showed a significant difference between brain regions (visual location: F=5.57, p < 10^-3^, auditory location: F=5.67, p < 10^-3^, and upcoming choice: F=11.6, p < 10^-9^). To compare differences between individual brain regions, we fitted a linear mixed effects model:

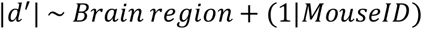

Here, |*d*’| is the absolute mean discrimination index across neurons, *Brain region* is a categorical fixed effect and *MouseID* is a random effect on the intercept (Figure S6f). To compare single neuron discriminability indices between naïve, trained mice and values obtained by shuffling the condition labels, we performed Welch’s t-test on the mean absolute discriminability index for each experimental session (Figure 5).

### Accumulator model

To investigate whether the structure of the sensory code in MOs can explain mouse behavior, we fed this code into an accumulator model (Figure 6, similar to a drift diffusion model(Ratcliff, 1978)). Since stimulus responses were sparse in MOs (140 auditory or visual location-selective neurons total from all experiments, as defined by the criteria of the previous section, i.e. 6%), we combined neural activity across all mice and experiments. To do so, we first obtained the PSTH for each stimulus condition, from –100 ms to 300 ms relative to stimulus onset. We then simulated 360 “trials” per stimulus condition by generating surrogate spike trains from a Poisson process with intensity given by these PSTHs. The stimulus conditions include unisensory visual trials with contrasts of 10, 20, 40, and 80%, unisensory auditory trials, and coherent and conflict audiovisual trials where the visual contrast is at 80%. This process yielded a time-dependent rate vector ***x***(*t*) for each trial, where *t* is time relative to stimulus onset.

The output of the accumulator model was a decision variable *d*(*t*), produced by linearly accumulating neural activity:

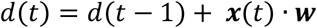

Here, ***w*** is a set of time-independent weights that were learned by the model to optimize the speed and accuracy of its responses but were not fit to mouse behavior. The choice of the model is defined by the sign of the decision variable when it crosses one of the thresholds: +1 or −1 for a rightward or leftward choice, and the reaction time of the model is the time of this threshold-crossing relative to stimulus onset (Figure 6d).

To learn the weight vector ***w***, we define a target decision variable *y* for each trial, set to 1 or −1 for rightward or leftward stimuli on unisensory and coherent trials. On conflict trials, where there is no correct response, the target decision variable is randomly set to 1 or −1 with equal probability.

The weights were learned by minimizing a loss function that compares the target decision variable with the model’s output decision variable for each trial:

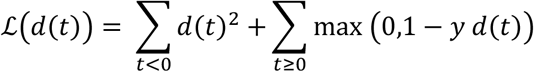

We used a mean-squared error loss before the stimulus onset (*t* < 0), to ensure that the model does not make a decision before the stimulus onset. After the stimulus onset (*t* ≥ 0) we use a hinge-loss error, which is zero when the decision variable is above the threshold for the correct choice and penalizes incorrect decisions and decision values below the decision threshold. The loss function was minimized with respect to the weights of the model via gradient descent using the ADAM optimizer (Kingma and Ba, 2014) with a learning rate of 0.01, and the gradient was obtained via automatic differentiation using the JAX library (Frostig et al., 2018). The model was trained on 70% of the trials for 300 epochs, and its behavior was evaluated on the remaining 30% of the trials (Figure 6e-h). To simulate the inactivation of the visual cortex in the right-hemisphere, we took the same learned model, but instead provided input where the activity of neurons that were previously identified as visual-left preferring neurons were decreased by 60% (Figure 6g). During training, the decision boundaries were set to +1 and −1. To account for the choice bias that was observed in mice, we performed grid search on the decision boundary values after model training in order to minimize the mean-squared error between the choice probability observed in the mice and in the model averaged across all stimuli conditions (Figure 6e). Decision boundaries were only fit on trials without simulated inactivation.

To simulate the inactivation of right MOs, we reduced the activity of right-hemisphere neurons by 60%. This manipulation did not recapitulate the lateralized effect of MOs inactivation (Figure S7j), because MOs neurons preferring either direction of stimulus are found equally in both hemispheres. To ask whether intra-hemispheric connections onto a downstream lateralized decision circuit could explain the lateralized effects of MOs inactivation, we trained another accumulator model with weights from neurons in the left and right hemisphere constrained to be positive and negative. This model was able to predict the lateralized effect of MOs inactivation (Figure 6h). To test whether this weight-constrained model recapitulated the lateralized effect of MOs inactivation better than the original accumulator model, we repeated the sampling and fitting procedure (as described earlier) 100 times for each model and performed a two-sample unpaired t-test on the mean-squared error between the model’s prediction of the log-odds and the observed log-odds from mouse behavior (p < 0.01).

To test whether sensory code in the MOs of naïve mice can produce the same behavior through the accumulator model (Figure 6c-d,f), we first subsampled the neurons recorded in the MOs of naïve mice so that the total number of neurons matched the number recorded in trained mice. To select the auditory and visual spatial neurons to be used in the accumulator model, we used the same shuffling procedure as above (see methods section on quantifying single neuron discrimination time). However, we adjusted the threshold which defines statistical significance so that the number of neurons selected from naïve mice to feed into the accumulator model matches that used from trained mice. Once neurons were selected, we fit the accumulator model with the same procedure used for trained mice (Figure 6e). This subsampling procedure was repeated 5 times, and the mean result across repeated subsamples was used for visualization (Figure 6c-d,f). To test whether the accumulator model fits (Figure 6e-h) are better than expected by chance, we compared the mean-squared error between the model’s prediction of the log odds log(*p*(*R*)/*p*(*L*)), compared to a null distribution obtained by fitting the same model after shuffling the stimulus conditions of each trial 100 times.

## Code & Data availability

The code used in the current study is available from the corresponding author on reasonable request. The datasets generated and/or analyzed during the current study are available from the corresponding author on reasonable request.

## Appendix 1

Here we show that optimally combining information from two inputs with independent noise requires the log odds be an additive function of the two inputs. This is a classical result of probability theory, whose significance to neuroscience has been discussed in several prior works (e.g. (Gold and Shadlen, 2002; Ma et al., 2006)). We provide a proof for the specific case of a binary left/right choice based on auditory and visual information.

Let *S* ∈ {*Ĺ, R*} represent the (left/right) location of the stimulus. A prior estimate for this location is captured by a prior probability distribution *p*(*S*). The two sensory inputs *A* and *V* follow conditional probability distributions *p(V* | *S*) and *p(A* | *S*). We assume them to be conditionally independent:

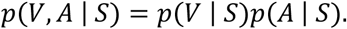

By Bayes’ theorem, the probability of the stimulus location given the inputs is:

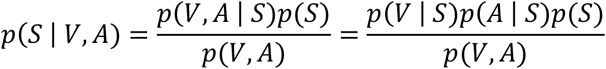

Write *p*(*R*) = *p*(*S* = *R*) and *p*(*L*) = *p*(*S* = *L*). Then, the log odds is given by:

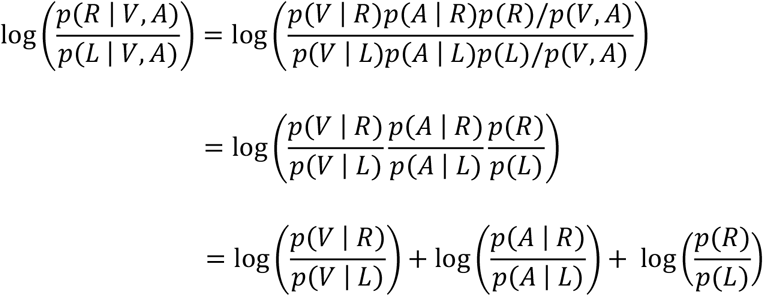

The first term depends only on *V*, the second only on *A*, while the third is a constant. Thus, the log odds is an additive function:

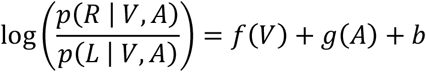

For a binary choice *p*(*L* | *V, A*) = 1 – *p*(*R* | *V*, *A*), so we can use the fact that the inverse function of 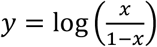 is the logistic function 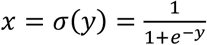, to obtain the formula

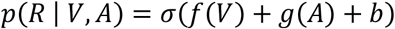

Thus, the assumption of independent noise in two sensory modalities implies that an optimal estimate of stimulus location uses the logistic function applied to an additive combination of the two modalities. While some psychophysical models instead use cumulative Gaussian models, the cumulative Gaussian function does not arise naturally in the same way.

In our task, the assumption that *p*(*V* | *S*) and *p*(*A* | *S*) are conditionally independent holds only approximately. The fact that mice combine evidence additively thus indicates they are following a heuristic strategy(Gardner, 2019). To demonstrate this, we compare the actual values of *p*(*V, A* | *S*) with those expected under the assumption of independence (Figure A1). To compute *p(V, A* | *S*) we use Bayes’ theorem: *p(V, A* | *S*) = *p*(*S|V*, *A*)*p*(*V, A*)/*p*(*S*). We define the stimulus location *S* to be the direction of wheel turn that will lead to reward on a particular trial. Thus, for unisensory or coherent multisensory stimuli on the right *p*(*R|V*, *A*) = 1, for conflict stimuli *p*(*R|V*, *A*) = 0.5, and for unisensory or coherent multisensory stimuli on the left, *p*(*R*|*V,A*) = 0. The prior probability *p*(*R*) = 0.5. Thus *p*(*V*, *A* | *R*) is the fraction of trials on which the combination (*V*, *A*) was presented if *V* and *A* are in conflict; twice this fraction if *V* and *A* are unisensory or coherent right; and 0 if *V* and *A* are unisensory or coherent left. These probabilities (Figure A1a) are not identical to those predicted by a conditional independence model *p*(*V* | *R*)*p*(*A* | *R*) obtained by multiplying the marginals of this distribution (Figure A1b).

**Figure A1.**
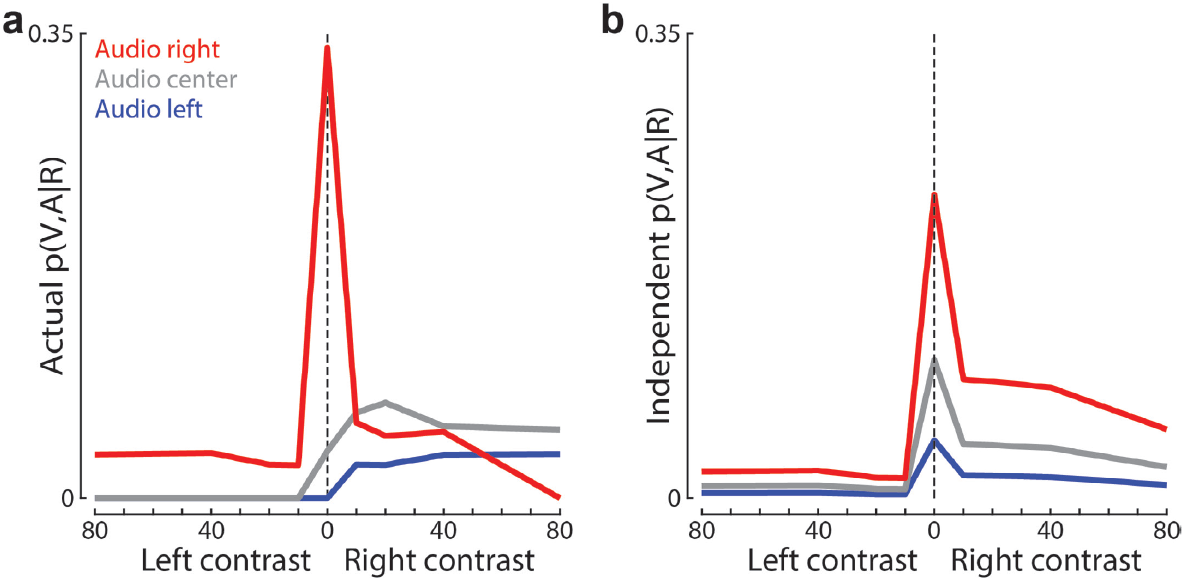
Conditional independence holds only approximately in our task. **(a)** The actual value of p(V, A | R) for each visual contrast when the auditory stimulus was presented on the right/center/left (red/grey/blue). Values were computed by Bayes’ theorem p(V, A |R) = p(R|V, A)p(V, A)/p(R), summing p(V, A) over ~156K trials from 17 mice. **(b)** As in (a), but for the conditional independence model p(V | R)p(A, | R).

**Figure S1.**
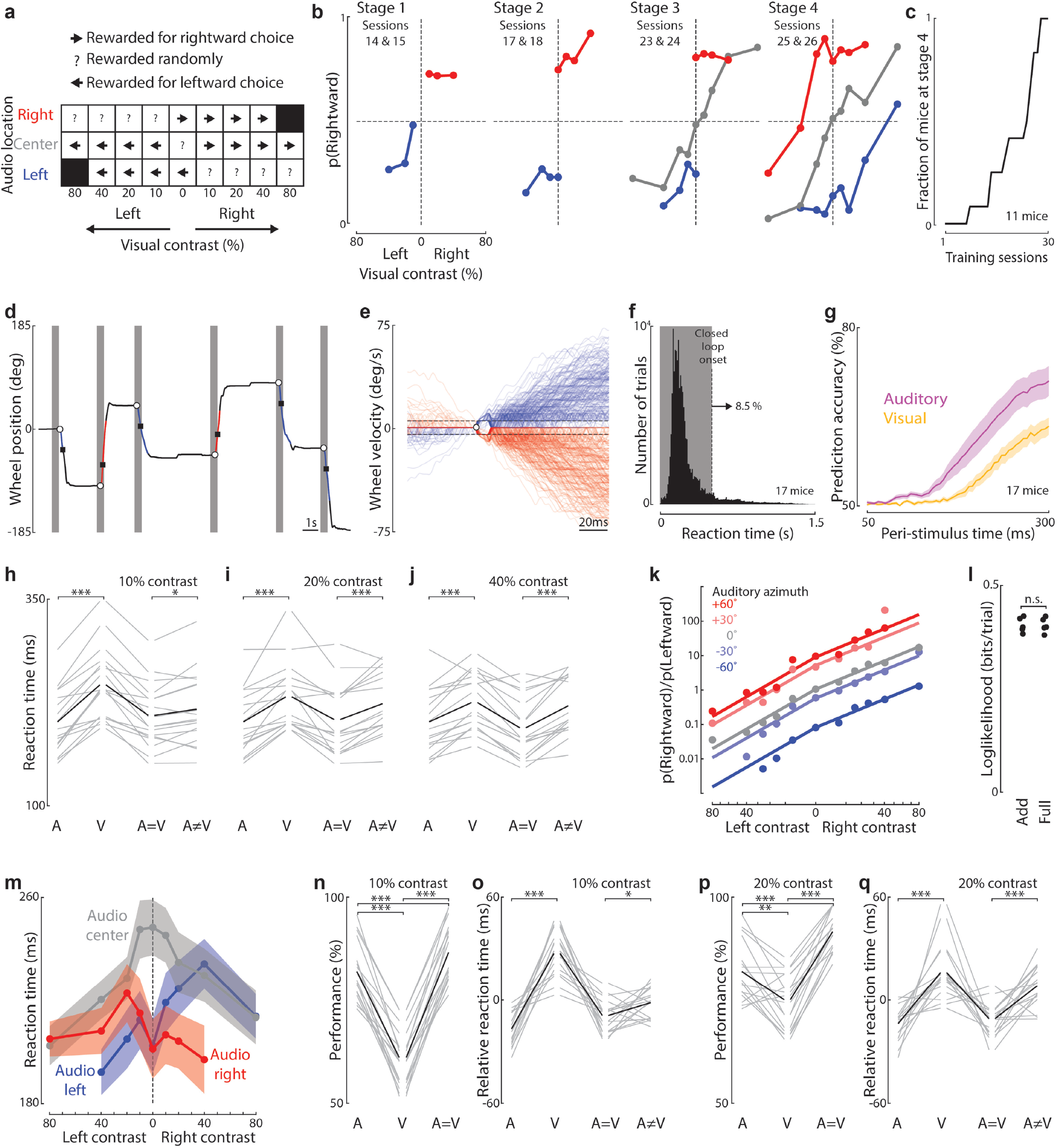
Training and behavioral classification. **(a)** Matrix of stimulus conditions and their corresponding reward criteria. **(b)** Training progression for one mouse. Mice first reach proficiency with audiovisual coherent trials (Stage 1), before we introduce unisensory auditory (Stage 2), unisensory visual (stage 3) and conflict (Stage 4) stimulus types. Plots for stages 1, 2, and 3, were made from the final two sessions of these stages. For stage 4, data is taken from the mouse’s first two sessions at that stage. **(c)** Number of sessions required to train mice. In our final training pipeline, ~80% of mice learned the task. Those that did learn required less than 30 sessions (median: 24.5). **(d)** Sample of wheel position trace. Grey regions indicate the 500ms open-loop period, beginning with stimulus onset. Open circles and squares indicate the movement onset and decision threshold (see Methods). **(e)** Zoom in to wheel velocity traces around the time of movement onset (open circle), for the same behavioral session as (d). Red and blue lines indicate leftward and rightward choices. Dashed lines represent the velocity thresholds for movement onset (see Methods). **(f)** Histogram of reaction times across 17 mice (~156K trials). Shaded region indicates the 500ms open-loop period, during which 91.5% of movements were initiated. **(g)** Accuracy of predicting whether a stimulus was presented on the left or right side from wheel velocity at each time point relative to stimulus onset, using a threshold obtained by minimizing an SVM loss function, for unisensory visual (yellow) or unisensory auditory (magenta) trials. Shaded areas indicate the standard error across 17 mice. Earlier predictions on auditory trials confirm that mice can decode the location of auditory stimuli earlier than visual stimuli (i.e. earlier auditory reaction times do not reflect guesses). **(h)** Comparing median reaction times for each stimulus type for each mouse, for trials with 10% visual contrast. Grey and black lines indicate individual mice and the mean across mice. ***: p < 0.001, *: p < 0.05 (17 mice, paired t-test); only comparisons between auditory and visual unisensory, and between conflict and coherent are shown. **(i)** As in (h), but for trials with 20% visual contrast. **(j)** As in (h), but for trials with 40% visual contrast. **(k)** Fit of the additive model to a mouse presented with 5 different auditory locations and 11 different contrast levels. Two additional parameters were used to fit the additional auditory stimuli (see Methods). **(l)** 5-fold cross-validated estimates of the loglikelihood for the additive and full (a parameter for each stimulus combination) behavioral models relative to a bias-only model. n.s. p > 0.05 **(m)** The mean (across 17 mice) of the median reaction times for each stimulus condition (~156K trials). Shading: standard error across mice. **(n)** Mouse performance (% rewarded trials) for unisensory auditory, unisensory visual, and coherent multisensory stimulus conditions (correct performance on conflict trials is undefined). This panel shows only trials with 10% visual contrast. Grey and black lines indicate individual mice and the mean across mice. ***: p < 0.001 (17 mice, paired t-test). **(o)** As in (h), but for reaction times relative to the mouse’s mean reaction time across all stimulus types. **(p)** As in (n), but for trials with 20% visual contrast. ***: p < 0.001 (17 mice, paired t-test). Data with 40% contrast is shown in Figure 1e. **(q)** As in (i), but with reaction times relative to the mouse’s mean reaction time across all stimulus types. Data with 40% contrast is shown in Figure 1b.

**Figure S2.**
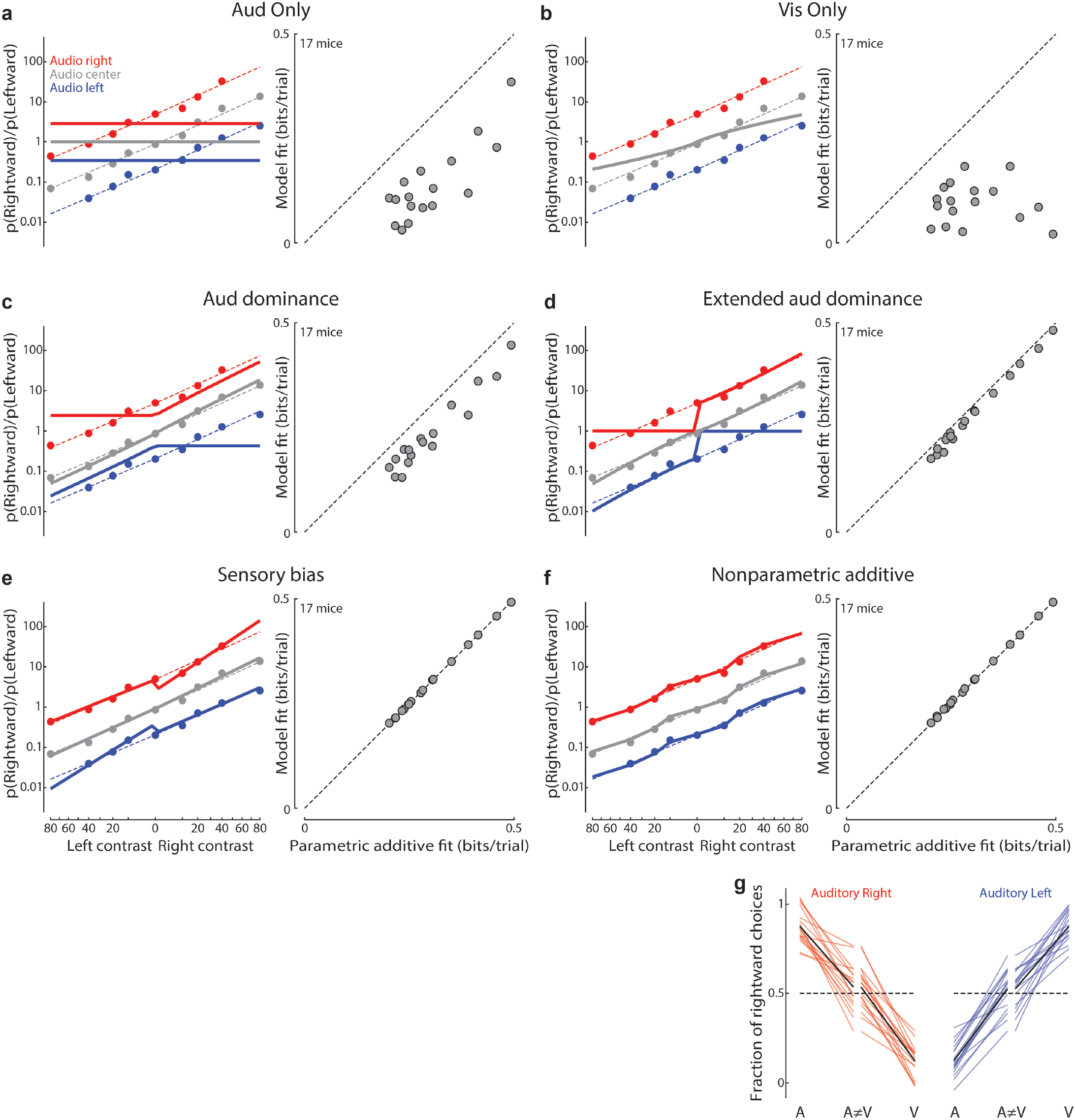
No evidence for sensory bias in behavior. **(a) Left**: Fit of an auditory-only model where both visual sensitivities (v_R_ and v_L_) are set to 0. Data plotted as the odds of choosing right vs. left (in log coordinates, Y-axis), as a function of visual contrast raised to the power γ. Closed circles: data combined across 17 mice. Red, grey, and blue curves: combined fit across all mice (17 mice ~156K trials). Dashed lines: fit of the additive model to the same data. The x-axis is visual contrast raised to the power γ from the additive model. **Right**: Log_2_-likelihood ratio for the auditory-only model versus the parametric additive model (both models assessed by 5-fold cross-validation relative to a bias-only model). The parametric additive model is a significantly better fit (p < 0.001, paired t-test). **(b)** As in (a), but for a visual-only model where both auditory sensitivities (a_R_ and a_L_) are set to zero. The parametric additive model is a significantly better fit (p < 0.001). **(c)** As in (a), but for a modified model that incorporates auditory dominance. Here, both visual sensitivities (v_R_ and v_L_) are set to zero on audiovisual conflict trials. The parametric additive model is a significantly better fit (p < 0.001). **(d)** As in (a) but for an extended auditory dominance model with an additional parameter to allow for a change in auditory sensitivity on conflict trials (see Methods). The parametric additive model is a significantly better fit (p < 0.001). **(e)** As in (a), but for a general sensory bias model with 4 additional parameters to allow for a change in visual or auditory sensitivity on conflict or coherent trials. The additive model is a subset of this model (see Methods). There is no significant difference in fit quality between this model and the parametric additive model (p > 0.05). **(f)** As in (a), but for the nonparametric additive model with a parameter for each auditory and visual stimulus condition. The superiority of the additive nonparametric model can be seen for example at visual contrast of +10, where the probability of a rightward choice is slightly below the parametric prediction for all three auditory conditions. The nonparametric additive model is a significantly better fit than the parametric additive model (p < 0.01). **(g)** Fraction of rightward choices for auditory, visual, and conflict stimulus types at the “matched” visual contrast for each mouse. Red or blue indicate stimulus conditions where the auditory stimulus was on the right or left. The “matched” contrast is the contrast that produces ~(1-X) fraction of rightward choices, where X is the fraction of rightward choices in the paired auditory stimulus condition. For both left and right auditory stimuli, the fraction of rightward choices on conflict trials is not significantly different from 0.5 (p > 0.05, t-test, n = 17 mice).

**Figure S3.**
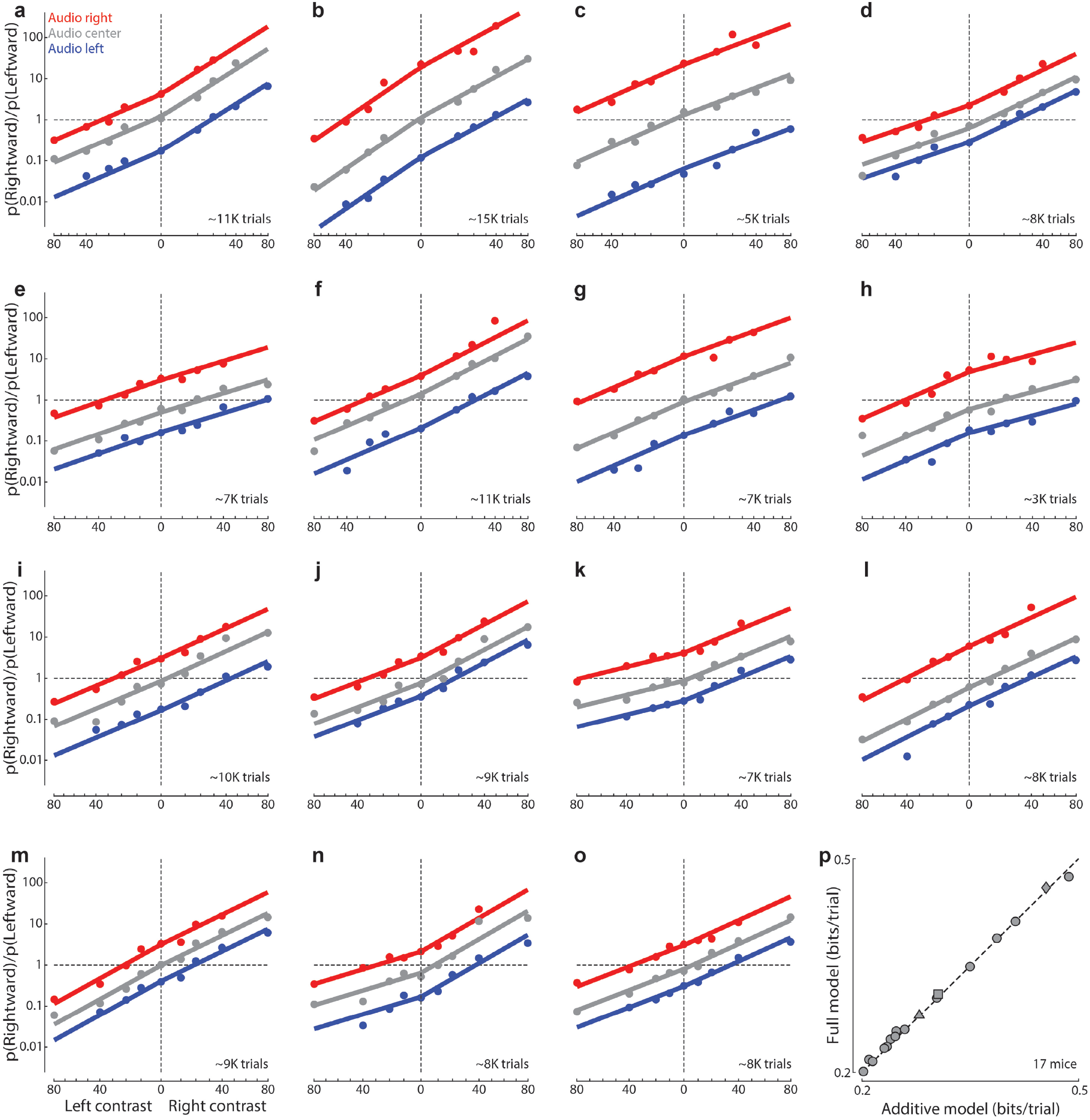
Additive model performance for each mouse. **(a-o)** Fits of the additive model to all 15 mice additional to those shown in Figure 1c, with different auditory and visual proficiencies, plotted as in Figure 1f. **(p)** Cross-validated loglikelihood ratio (relative to a bias-only model) for the full model versus the additive model trained on only unisensory stimulus conditions, plotted as in Figure 1h. There is no significant difference between models. p > 0.05 (17 mice, paired t-test).

**Figure S4.**
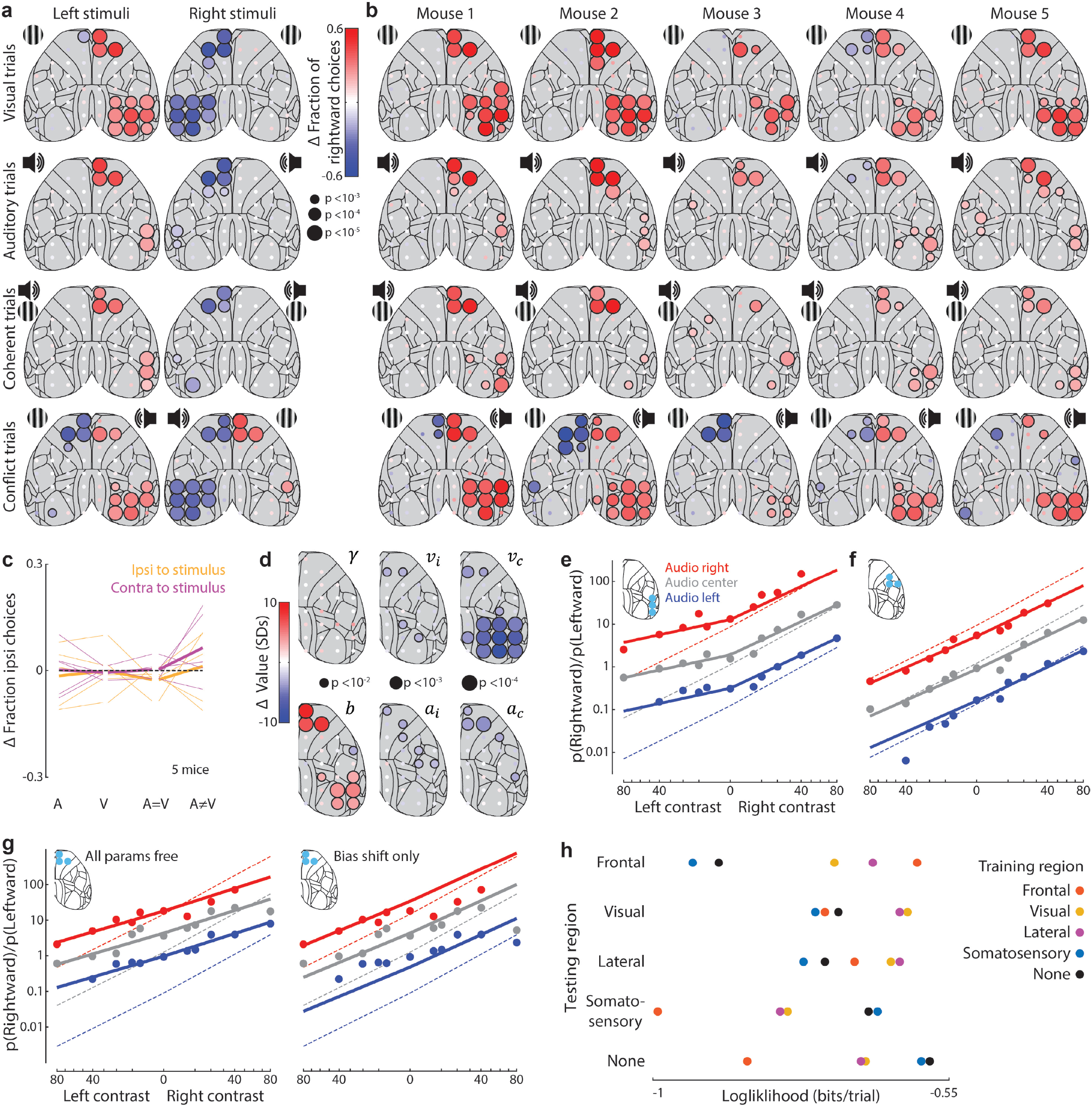
Further analysis of optogenetic inactivation effects on choice, model, and brain regions. **(a)** Change in the fraction of rightward choices for 52 stimulation sites for unisensory visual, unisensory auditory, coherent, and conflict stimulus conditions. The position of the grating and loudspeaker symbol indicate the locations of the visual and auditory stimuli in each case; Left and right columns represent mirrored cases of the same stimulus condition. Dot color indicates change in the fraction of rightward choices and dot size represents statistical significance (5 mice, permutation test, see Methods). **(b)** As in (a), but each column represents an individual mouse, and mirrored trials were combined, reversing if necessary, so that the stimulus (visual in conflict trials) was always on the left. Dot color indicates change in the fraction of rightward choices. **(c)** Change in the fraction of ipsilateral choices on trials when inactivation target was outside the brain (on the dental cement and/or skin of the mouse). No significant effect was observed for any stimulus condition for ipsilateral (gold) or contralateral (magenta) inactivation sites. p > 0.05 (paired t-test). **(d)** As in main text Figure 2f, but also allowing the contrast gain saturation parameter y to be fit. No significant changes in the contrast gain parameter (γ) were observed (5 mice, permutation test, see Methods). **(e)** Fit of additive model to trials in which any of 3 sites in lateral sensory cortex were inactivated, plotted as in Figure 2g-h. Trials with inactivation of left lateral sensory cortex were included in the average after reflection. The deviation in model parameters was significant (paired t-test, p < 0.05). **(f)** As in (e), but for trials when somatosensory cortex was inactivated (5 mice, 6689 trials). The deviation in model parameters was significant (p < 0.05). **(g)** Fit of the additive model to right frontal cortex inactivation trials, with all parameters free (left, as in Figure 2h) or with only the bias parameter free to change from its non-inactivation value (right). This “Bias shift only” approach resulted in a worse fit (paired t-test, p < 0.05), due to its inability to model changes in sensory sensitivity. **(h)** Loglikelihood for trials when each testing region was inactivated, evaluated using model parameters from a fit to trials in which the training region was inactivated. In each case, loglikelihood is highest when testing and training regions are the same. Every inter-region loglikelihood is significantly worse (p < 0.05, shuffle test, Bonferroni-corrected).

**Figure S5.**
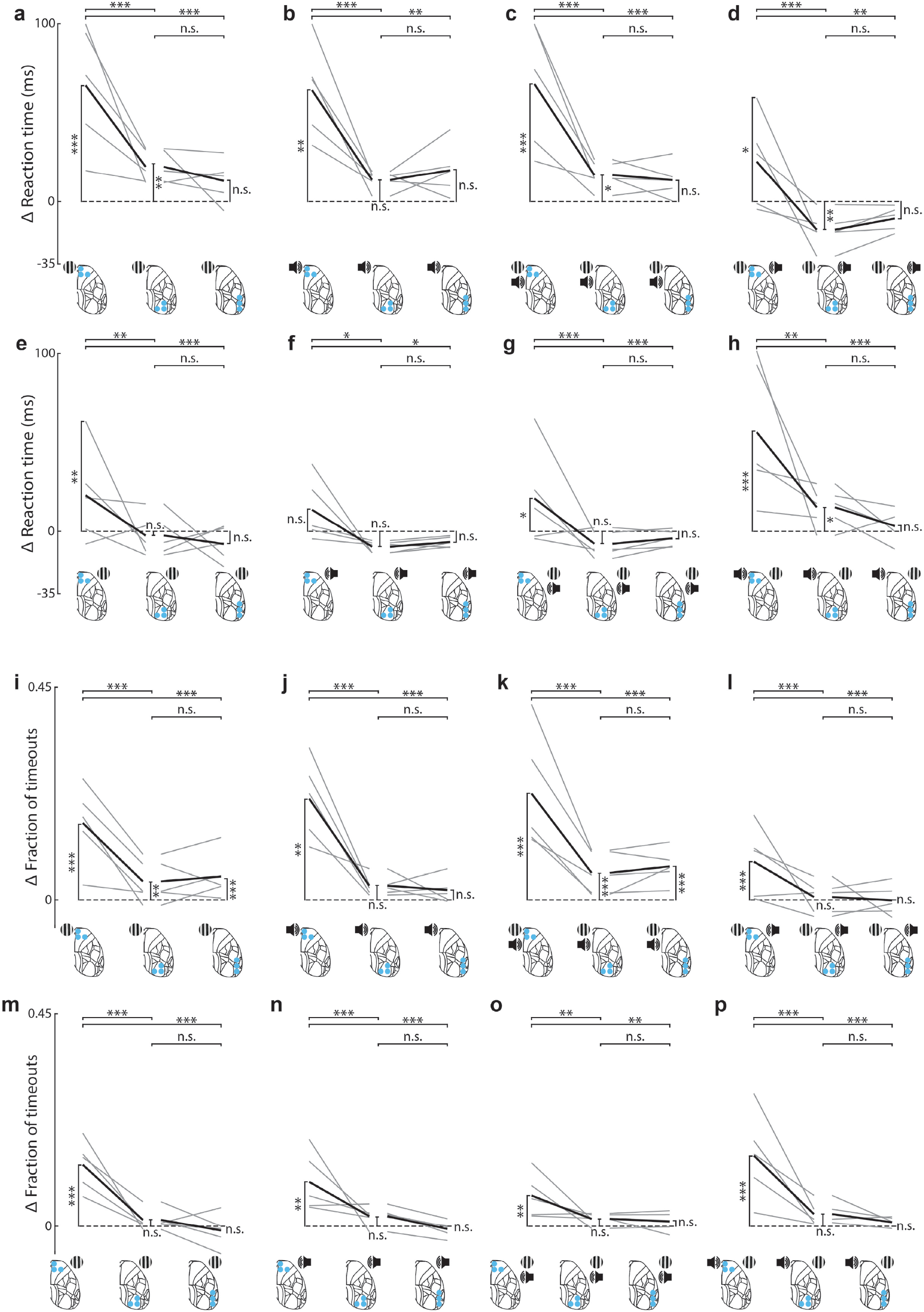
Effects of inactivation on reaction time and fraction of timeouts. **(a)** Change in reaction times when frontal, visual, and lateral sensory cortex were inactivated (left, center and right) on visual trials, relative to non-inactivated trials with the same (contrast-matched) stimuli. Inactivated regions could be on the left or right, but were always contralateral to the visual stimulus. Grey and black lines indicate individual mice (n = 5) and the mean across mice. The values for each mouse were calculated by first computing the median reaction time for each contrast, then taking the mean of these over contrasts. Values above 100 ms were truncated to 100 ms for visualization but not analyses. n.s: p > 0.05, *: p < 0.05, **: p < 0.01, ***: p < 0.001 (linear mixed effects model). **(b)** As in (a), but for auditory trials. **(c)** As in (a) but for coherent trials. **(d)** As in (a), but for conflict trials where the visual stimulus was contralateral to the inactivated region. **(e-h)** As in (a-d) but for trials of reversed laterality such that the stimulus (visual in the case of conflict trials) was ipsilateral to the inactivated region. **(i-p)** As in (a-h) but for the change in the fraction of timeout trials (no response within 1.5s) rather than reaction times.

**Figure S6.**
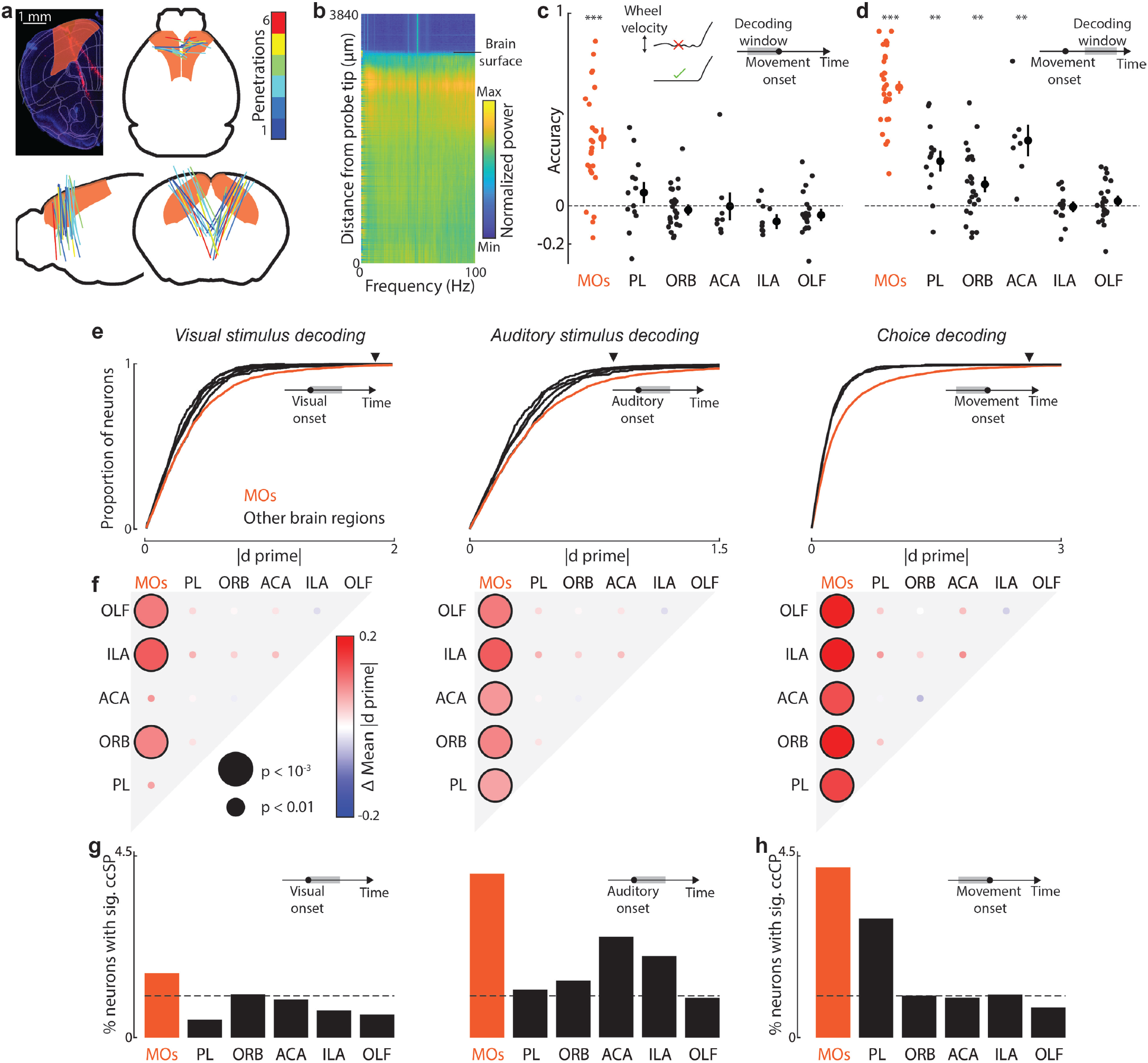
Electrophysiology methods and controls. **(a)** Histological reconstruction of electrode tracks. Top left: Example coronal section (DAPI-staining, blue) at 1.86mm anterior to bregma with tracks from 2 insertions. Slices were registered to the Allen Atlas (Wang et al., 2020) (overlaid, with MOs highlighted in orange) using custom software (see Methods). Right and bottom: reconstructed trajectories of all recordings analyzed, projected onto coronal, sagittal, and horizontal planes. Color indicates the number of penetrations along a given trajectory (one DiI-labelled penetration, same angles and point of insertion). **(b)** Normalized power spectrum (log(power), z-scored across probe depth) versus depth along probe for a single penetration. The point where the probe enters the brain can be identified by a sudden increase in power. **(c)** Cross-validated accuracy (relative to a bias model, see Methods) of an SVM decoder trained to predict rightward or leftward choices based on population spiking activity time-averaged over a time window 0 ms to 130 ms preceding movement. Trials where mice made sub-threshold movements prior to movement onset were excluded (illustrated in inset). Each point represents the decoding accuracy from neurons in one brain region (Secondary motor (MOs), orbitofrontal (ORB), anterior cingulate (ACA), prelimbic (PL), infralimbic (ILA), or Olfactory (OLF)), from a single experimental session. ***: p < 0.001. **(d)** As in (c), but with a time window of 150 ms to 300 ms after movement onset (see Methods). Trials where mice made sub-threshold movements were not excluded. **: p < 0.01, ***: p < 0.001. **(e)** Cumulative values of the discriminability indices (absolute d-prime, see Methods) across sessions. Lines represent MOs (orange) and all other recorded brain regions (black). Arrows indicate the absolute d-prime values for the example neurons in (Figure 3b-d). **(f)** Significance for inter-region comparison of d-prime values from (e). d-prime values in MOs were significantly greater than all other regions for encoding auditory location and upcoming choice, but not significantly greater than ILA or ACA when considering visual location (Linear mixed effects model, see Methods). **(g)** Proportion of neurons sensitive to visual (left) or auditory (right) stimulus location, estimated with combined conditions stimulus probability analysis (ccSP; see Methods), after controlling for the other stimulus and for choice, using neural activity time-averaged over a window 0 to 300 ms after stimulus onset. In both cases, MOs has the highest proportion of significant neurons (more than 700 neurons were recorded in each area). Dashed line indicates the percentage of neurons expected by chance. **(h)** As in (g), but using combined conditions choice probability (ccCP; see Methods) to estimate the proportion of neurons sensitive to the upcoming choice, after controlling for both stimuli, using neural activity time-averaged over a window 0 to 130 ms before movement onset.

**Figure S7.**
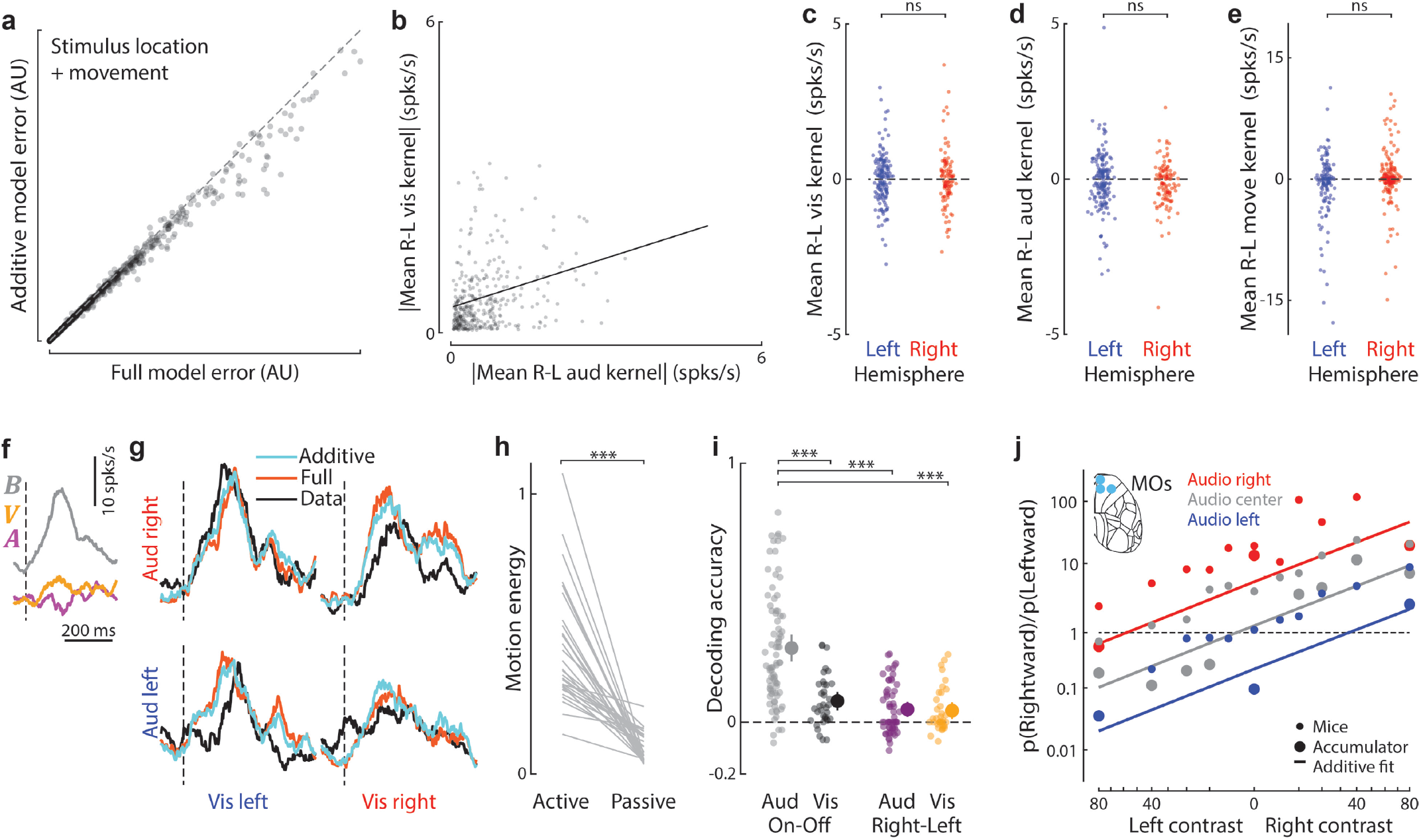
Additional analysis for recordings during behavior. **(a)** Prediction error for each neuron for the additive and full sensory-movement models (see Methods). The full model has significantly higher error. (p < 0.01, 2183 cells, linear mixed-effects model). For visualization purposes, the largest 1% of errors were excluded from the plot, but not from statistical analysis. **(b)** Absolute value of time-averaged visual R-L kernel (V), versus absolute value of time-averaged auditory R-L kernel (A), after fitting the additive model under passive conditions for significant neurons. A time window from 0 to 300 ms after stimulus onset was used in both cases. Absolute responses are correlated, suggesting that if a neuron is predictable from spatial stimuli in one modality, it is more likely to be predictable from the other modality (p < 10^-5^, 2509 cells, linear mixed-effects model). **(c)** Signed time average of the visual R-L kernel (V, Eqn 5, 0 ms to 300 ms after stimulus onset) versus recorded hemisphere, after fitting the additive neural model (N = 0) under passive conditions (D, M = 0) (see Methods). There is no significant lateralization in spatial preference (p > 0.05; 273 cells, linear mixed-effects model). **(d)** As in (c), but for the auditory R-L kernel (A). (p > 0.05; 287 cells). **(e)** As in (c), but the movement R-L kernel (D, −200 to 400 ms relative to movement onset), after fitting the additive model during active behavior (p > 0.05; 209 cells). **(f)** Example sensory kernels from fitting the additive neural model to a single neuron under passive conditions. The selected neuron has opposing sensitivities for auditory (left-preference) and visual (right-preference) stimulus locations. B: baseline kernel; V: visual direction kernel; A: auditory direction kernel. **(g)** Cross-validated fit using the model kernels from (f) to neural responses under passive conditions for all audiovisual combinations. Cyan and orange lines show predictions of the additive and full models, black line shows test-set average responses. **(h)** Difference in facial video motion energy (see Methods) when mice are performing the behavior (active) versus passive presentation of stimuli (passive). Mice exhibit significantly less motion under passive conditions. *: p < 0.001 (30 sessions, paired t-test). **(i)** Neurons were tested individually to see if they encoded the presence or location of visual and auditory stimuli (see Methods). The plot shows the single-neuron decoding accuracy of all significant neurons for each discrimination. Neurons encoding auditory stimulus presence (grey, 82 cells) have higher decoding accuracy than all other categories (36/59/36 cells for black/magenta/gold). *: p < 0.001 (t-test). **(j)** Mean behaviour of the accumulator model with the activity of all neurons in the right hemisphere reduced by 60% to simulate inactivation of right MOs (large circles), plotted with the mean behaviour from MOs-inactivated mice (5 mice, small circles, cf. Figure 2h). Solid lines represent the fit of the additive model to the accumulator model output. This model cannot capture the rightward choice bias following right MOs inactivation, because MOs neurons preferring either direction of stimulus are found equally in both hemispheres.

